# Sequence and annotation of 42 cannabis genomes reveals extensive copy number variation in cannabinoid synthesis and pathogen resistance genes

**DOI:** 10.1101/2020.01.03.894428

**Authors:** Kevin J. McKernan, Yvonne Helbert, Liam T. Kane, Heather Ebling, Lei Zhang, Biao Liu, Zachary Eaton, Stephen McLaughlin, Sarah Kingan, Primo Baybayan, Gregory Concepcion, Mark Jordan, Alberto Riva, William Barbazuk, Timothy Harkins

**Affiliations:** Medicinal Genomics, 100 Cummings Center, suite 406-L, Beverly, MA 01915; The University of Florida Interdisciplinary Center for Biotechnology Research, Gainesville, Florida 32611; Pacific Biosciences, 1305 O’Brien Dr, Menlo Park, CA 93025; Minnibis, 1708 17th Ave, Longmont, CO 80501

**Author notes:** Corresponding Author: Kevin McKernan, Medicinal Genomics, 100 Cummings Center, suite 406-L, Beverly, MA 01915.

**Keywords:** Cannabis, Chitinases, Thaumatin-Like Protein, Cannabinoid synthases, single molecule sequencing, whole genome assembly, copy number variation.

## Abstract

Cannabis is a diverse and polymorphic species. To better understand cannabinoid synthesis inheritance and its impact on pathogen resistance, we shotgun sequenced and assembled a *Cannabis* trio (sibling pair and their offspring) utilizing long read single molecule sequencing. This resulted in the most contiguous *Cannabis sativa* assemblies to date. These reference assemblies were further annotated with full-length male and female mRNA sequencing (Iso-Seq) to help inform isoform complexity, gene model predictions and identification of the Y chromosome. To further annotate the genetic diversity in the species, 40 male, female, and monoecious cannabis and hemp varietals were evaluated for copy number variation (CNV) and RNA expression. This identified multiple CNVs governing cannabinoid expression and 82 genes associated with resistance to *Golovinomyces chicoracearum*, the causal agent of powdery mildew in cannabis. Results indicated that breeding for plants with low tetrahydrocannabinolic acid (THCA) concentrations may result in deletion of pathogen resistance genes. Low THCA cultivars also have a polymorphism every 51 bases while dispensary grade high THCA cannabis exhibited a variant every 73 bases. A refined genetic map of the variation in cannabis can guide more stable and directed breeding efforts for desired chemotypes and pathogen-resistant cultivars.

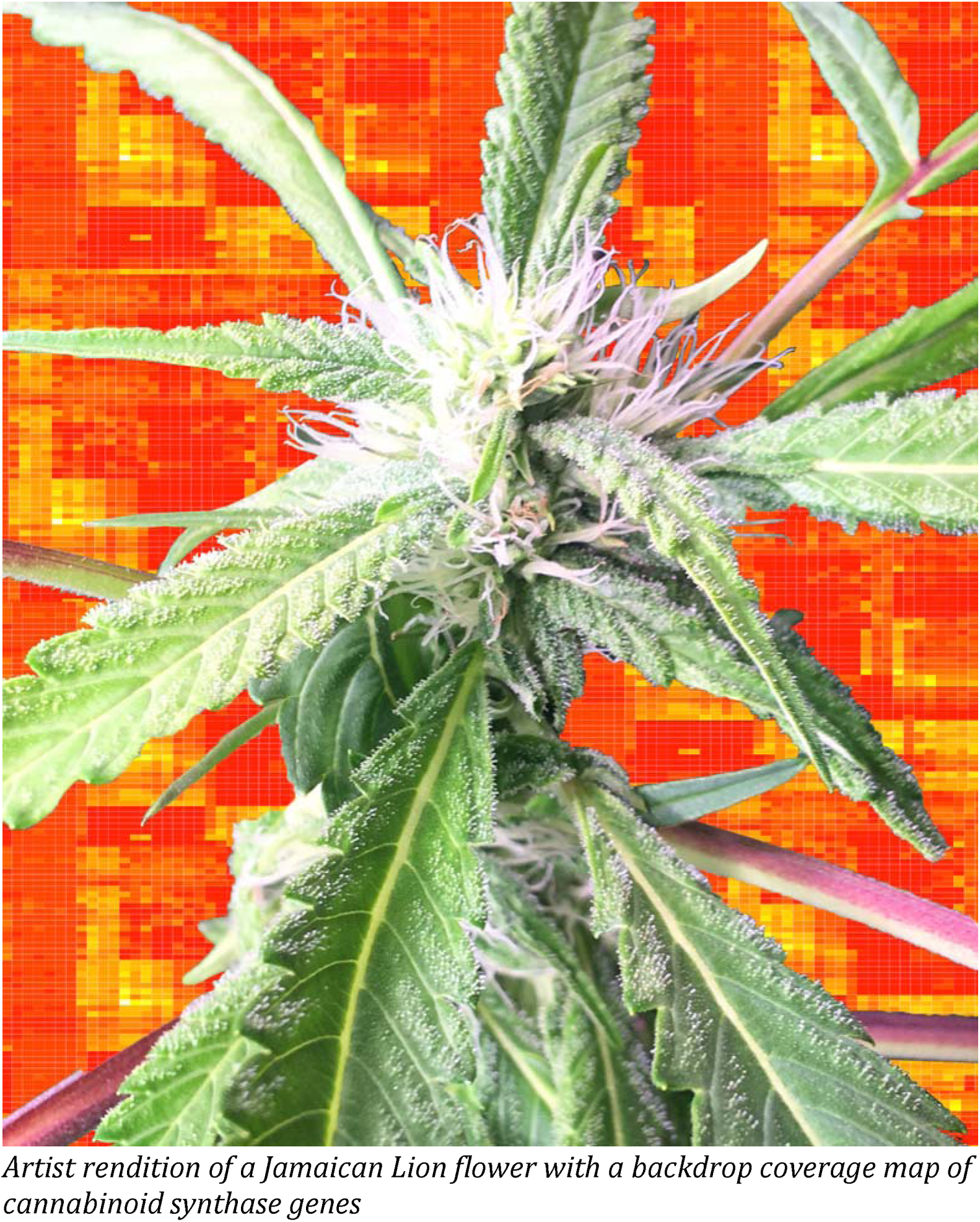
Sequence and annotation of 42 cannabis genomes reveals extensive copy number variation in cannabinoid synthesis and pathogen resistance genes

## Introduction

*Cannabis sativa* L. is one of the earliest domesticated plants (Ren et al. 2019). Classified independently by Linnaeus and Lamarck, hemp fiber was used by Marco Polo and James Cook for rope, sails, paper, and ship caulk (Clarke and Merlin 2013). The extensive maritime use of cannabis has played a role in its spread around the globe, creating an interesting admixture of landrace genetics. The suitability of the common subspecies vernacular of *Cannabis sativa*, subsp. *sativa indica*, and *ruderalis* has been hotly debated but infrequently verified with genomic surveys. Segregation of fiber-based cannabis (hemp) and drug type cannabis (sometimes referred to as marijuana) has been genetically resolved (Small; Hillig and Mahlberg 2004; Fischedick et al. 2010; van Bakel et al. 2011; Hazekamp and Fischedick 2012; Sawler et al. 2015; Ryan C. Lynch 2016; Soorni et al. 2017). Hemp genetics are ancestral and often produce distinct male and female flowers on a single plant (monoecious). Dioecious hemp and cannabis can be hermaphrodites but this usually entails bisexual flowers that are distinct from flowers found on monoecious phenotypes. Drug type cannabis has undergone selection for female flowers that produce high levels of tetrahydrocannabinolic acid (THCA). This selection has produced predominantly dioecious female varieties for modern cannabinoid production (Grassa et al. 2018; Laverty et al. 2019).

Cannabis is diploid and has 10 chromosomes (2n=20) and an XY sex chromosome system (Divashuk et al. 2014). While dioecious genetics are preferred for cannabinoid production, hermaphroditic traits still circulate in the drug type varietals, complicating mass production of cannabinoids. Since pollinated female flowers produce lower levels of cannabinoids and terpenes, male drug type plants are visually or genetically tested and culled from grows to prevent pollination of female plants. Visual sexual differentiation occurs in the midlife cycle of the plant, while genetic screening can eliminate males earlier to conserve expensive indoor growing real estate. Female plants with hermaphroditic tendencies are more difficult to detect and remove using visual or genetic screening methods.

To avoid the propagation of Y chromosomes in naturally-crossed cannabis seeds, indoor growers often resort to tissue culture or cloning of female mother plants. This can lead to monocultures and increased risks associated with pathogen exposure (Wally and Punja 2010). Another approach more common in outdoor cannabidiolic acid (CBDA) production utilizes the induction of hermaphroditism with silver nitrate or silver thiosulfate and ethephon treatment (an ethylene blockers and ethylene mimetic respectively). These phytohormone modulators reverse the sex phenotype of some female and male varietals respectively. Only sex reversal of female plants with ethylene blockers results in pure XX pollen production (Mohan Ram and Sett 1982). Application of XX pollen to female flowers results in ‘feminized’ seeds but can also increase the incidence of plants with hermaphroditic capacity. The hermaphroditic capacity of varietals is believed to be a heritable trait but this trait has yet to be mapped to any genomic coordinates. Sex chromosomes in flowering plants usually evolve from autosomes (Harkess et al. 2016; Harkess et al. 2017). Likewise, modern monoecious hemp varietals tend to decay into dioecious varietals with inbreeding but little is known about their chromosomal structures (Clarke 2017).

Cannabinoid and terpene production by plants is linked to both attraction of pollinators and responses to plant pathogens (Penuelas et al. 2014; Andre et al. 2016; Allen et al. 2019; Lyu et al. 2019). Cannabis plants are classified into 5 categories reflecting their THCA, CBDA, and CBGA expression. Type I plants express predominantly THCA. Type II plants express a near equal mixture of THCA and CBDA. Type III plants express predominantly CBDA, while Type IV plants express neither CBDA nor THCA and predominantly express the CBGA precursor. Type V plants express no cannabinoids. Despite successful breeding efforts to deliver higher cannabinoid and terpene content, plant pathogens are still a significant contributor to crop loss in cannabis production due to the lack of disease-resistant varieties (Kusari 2013; Backer et al. 2019). Many jurisdictions mandate cannabis microbial safety testing targeting epiphytic and endophytic plant pathogens that have been clinically linked to aspergillosis in immuno-compromised cannabis patients (McKernan et al. 2015; Remington et al. 2015; McKernan et al. 2016).

To better understand cannabis sex evolution, cannabinoid expression and pathogen resistance, we sequenced and assembled male and female cannabis genomes (cultivar ‘Jamaican Lion mother’ and ‘Jamaican Lion father’) with their offspring (JL1 – JL6) to identify the Y chromosome. We further annotated these references with full-length mRNA (Iso-Seq) sequencing of 5 tissues (female flowers, female seeded flowers, male flowers, female leaves and female roots) and *in silico* gene model predictions using MAKER2 (Cantarel et al. 2008; Campbell et al. 2014a; Campbell et al. 2014b; Law et al. 2015). To confirm the 118Mb list of putative Y contigs, we utilized Illumina’s NovaSeq platform to whole genome sequence 40 hemp and drug-type cultivars. Using the coverage maps of 9 male genomes, we were able to confirm putative Y contigs and male-specific genes (Prentout 2019). We further classified these variants to identify highly damaging mutations in protein coding regions of the genome. Additionally, these whole genome sequence data were utilized to assess copy number variation (CNV) in critical genes in the terpene synthase pathway, cannabinoid synthase pathway and the pathogen response pathway (Pisupati et al. 2018). This refined map of protein coding variants and copy number variants can facilitate directed breeding efforts for desired chemotypes, pathogen resistance and a better understanding of rare cannabinoid synthesis (Citti 2019).

## Results and Discussion

### Sequencing, assembly and annotation

Two maternal draft assemblies were released previously with 652Kb N50s and 3.8Mb N50 contig sizes respectively (McKernan 2018a). To complement these assemblies, a male sibling and F1 female offspring were also sequenced and assembled. FALCON-Unzip was used to assemble over 85 Gb of sequence for each Trio member (Chin et al. 2016). These data were polished with Arrow and annotated with Iso-Seq data and MAKER2. This delivered 3 assemblies with N50s over 1.6Mb and benchmarking single copy orthologs (BUSCO) scores >96% (Table 1, Supplemental Fig S1.). Multiple tools were explored to remove duplicate BUSCO genes with Polar star and Purge haplotigs (Roach et al. 2018). Any condition that reduced the completeness score and showed signs of purging cannabinoid synthase genes was rejected. Haplotig purging was performed only on the maternal reference using default conditions with Polar star and Purge haplotigs but BUSCO duplicate scores remain at 12.6%. In polymorphic genomes with known CNVs in cannabinoid synthase genes, haplotig contamination can be hard to discern from real copy number variation. The assumption of genome-wide diploidy can be misleading, with elevated structural variation and evidence of long terminal repeat (LTR)-driven gene expansions seen in Laverty *et al.* (Laverty et al. 2019). If a long-read assembler (such as FALCON) finds contigs divergent enough to split them in the primary assembly, we opted to leave those in the reference. Alternative haplotigs discovered in the unzipping and phasing process were omitted from the CNV analysis as not all parts of the genome were divergent enough between the related maternal and paternal lines to be unzipped and phased. Inclusion of alternative haplotigs created false hemizygous deletions when comparing the genome coverage to highly similar and collapsed paternal and maternal contigs.

This study is one of the first to utilize Sequel II CLR reads and CCS reads to assemble and annotate a cannabis genome and represents the first type II cannabis genome assembled containing both functional THCAS and CBDAS alleles. As a result, many reads are longer than previously reported N50 contig lengths in cannabis genomes. CCS sequencing on RNA molecules generates over 30× coverage of individual molecules resulting in multi-kilobase reads with over Q30 accuracy. This is helpful to identify accurate splice junctions and transcript isoforms. The read lengths are often longer than nuclear NUPTs or NUMTs enabling more confident heteroplasmy detection in organellar genomes. These reads also enable structural variation detection critical to resolving the CNVs defining the most common Bt:Bd allele observed in most cannabis and hemp cultivars. This is the first genome-wide population survey conducted on a reference-grade genome and the longer reads greatly simplify the analysis of gene family expansions in cannabinoid and terpene synthase gene families.

With these data, we identified male- and female-specific contigs with whole genome alignments from 40 Illumina sequenced cultivars (Figure 1). The parental lines and 6 F1 offspring are included as controls in the Illumina sequenced 40 genomes (Figure 2).

**Figure 1:**
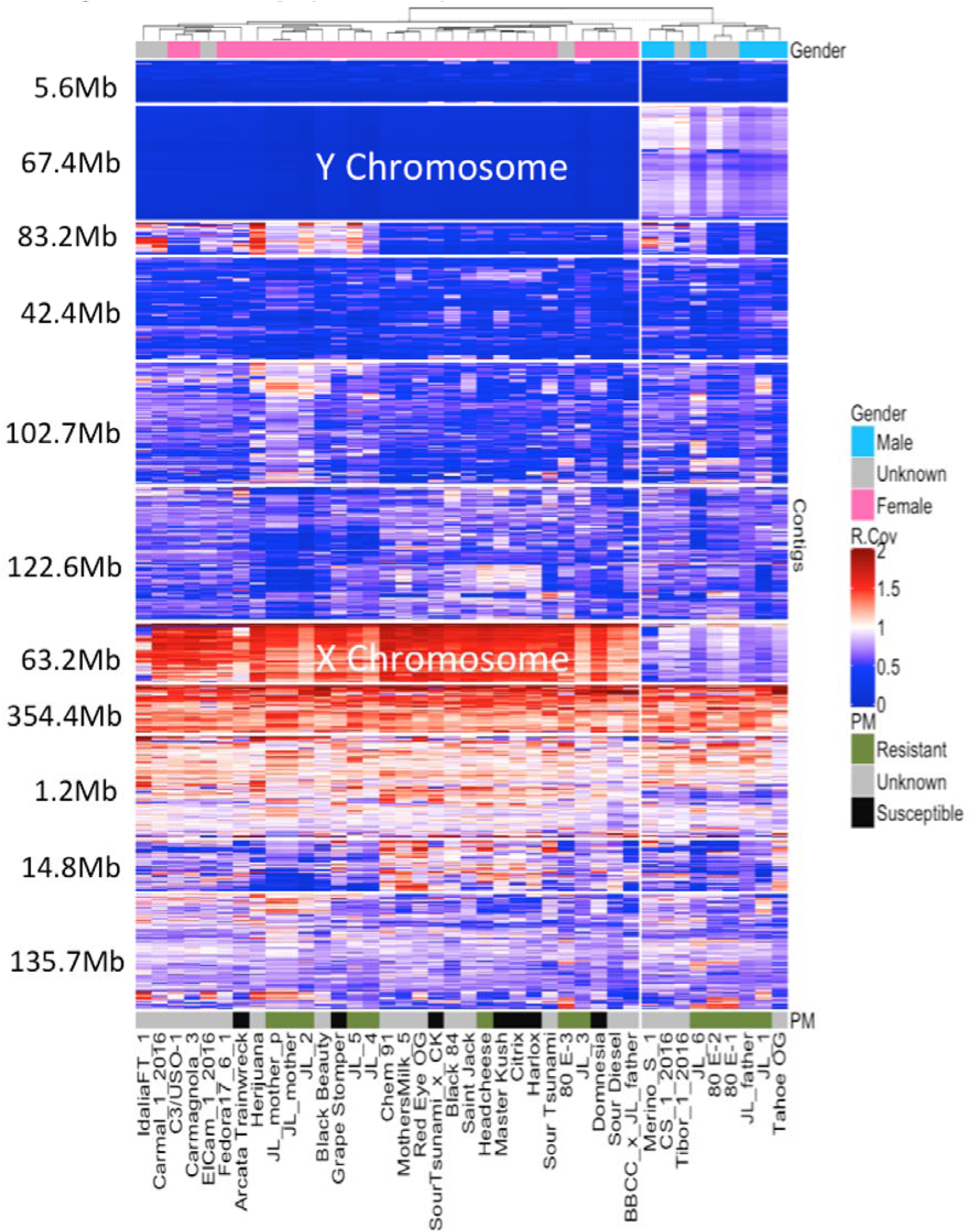
Coverage maps of Illumina sequence across the paternal genome. Paternal genome was not unzipped due to the presence of the pseudoautosomal region of the Y chromosome. Relative coverage of Illumina reads across 1,298 contigs (y-axis) for 40 genomes (x-axis). The nine male genomes are to the right. Female genomes on the left demonstrate no coverage (Blue) over male specific contigs and double the coverage (Red) over X contigs. Some cultivars were provided with unknown gender represented in grey on the bottom x-axis. The contigs are organized into 11 clusters based on their coverage correlation but do not always represent chromosomes. Cumulative length of the contig clusters are displayed on the y-axis.

**Figure 2A.**
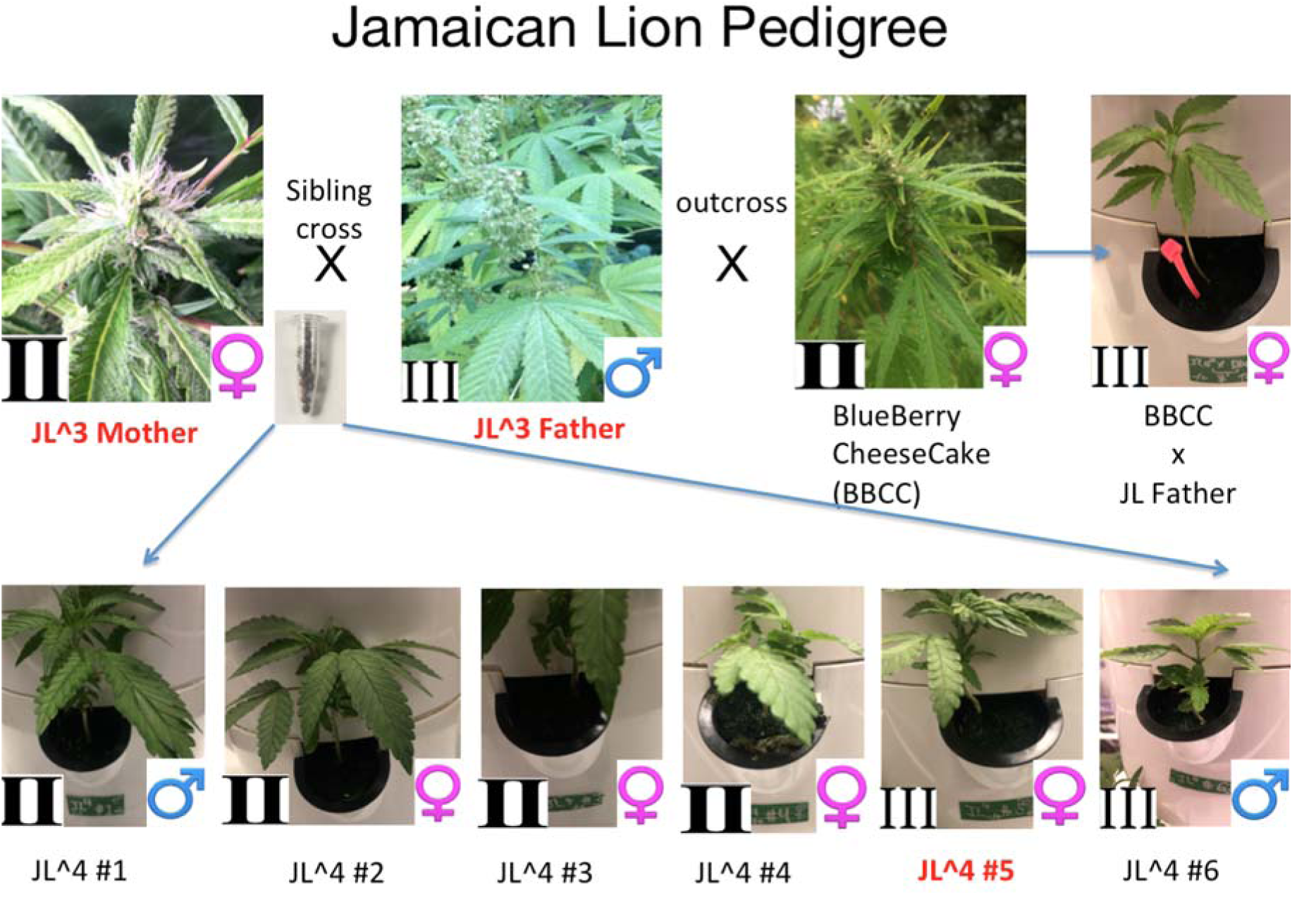
The Jamaican Lion pedigree. All samples were whole genome sequenced with Illumina NovaSeq® platform. Samples with names in Red were also sequenced and assembled with Pacific Biosciences Sequel II platform.

**Figure 2B.**
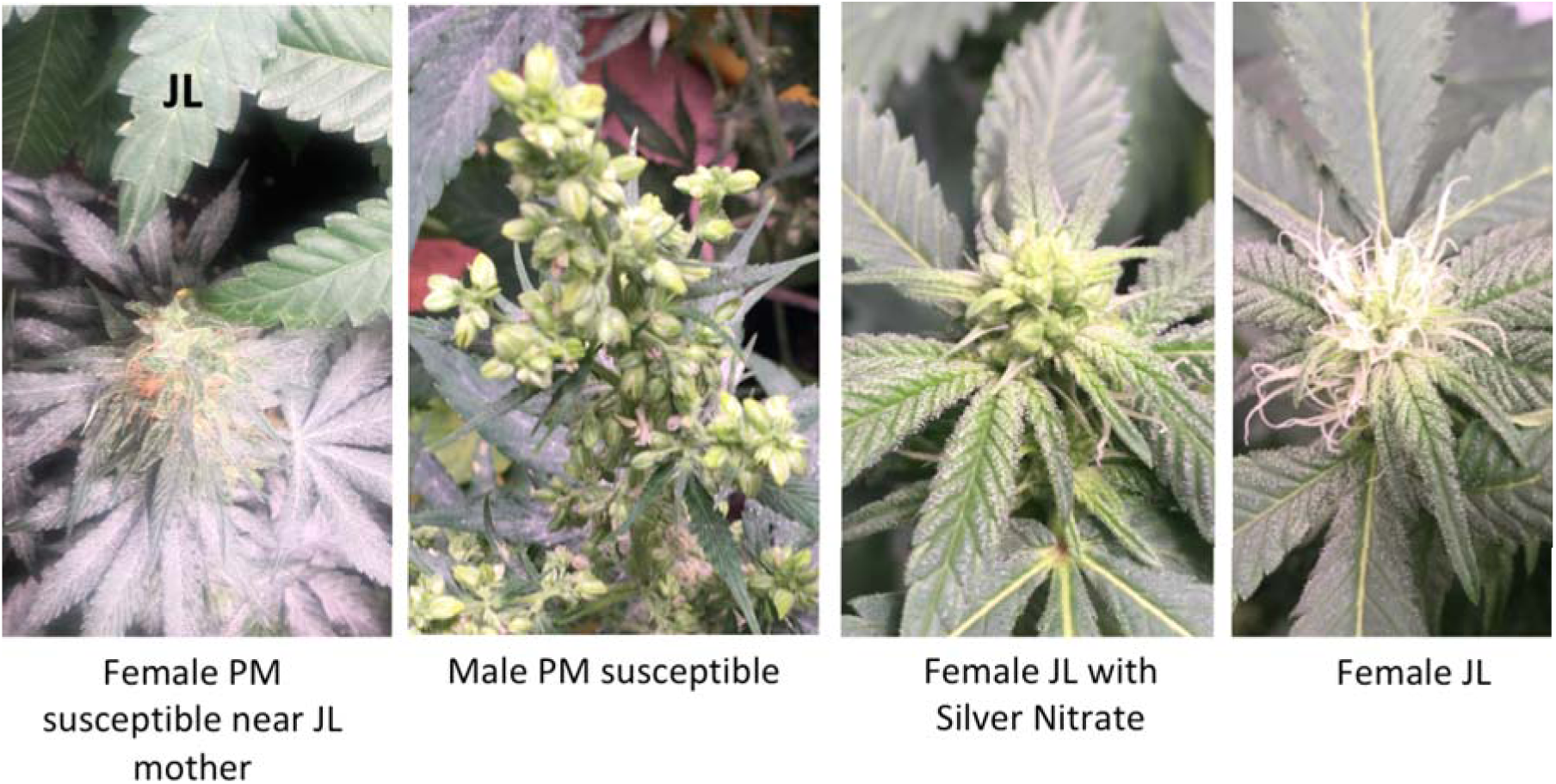
Left) Attempted inoculation of Jamaican Lion mother clone with *G. chicoracearum*. Middle-left) Male *G. chicoracearum*-susceptible variety. Middle-right) Jamaican Lion mother clone treated with silver nitrate to induce hermaphroditic female flowers. Right) Non-treated Jamaican Lion mother flower(XX).

### Identification of the Y chromosome

To identify the Y chromosome, 40 genomes were aligned to the paternal Pacific Biosciences reference assembly. Nine male genomes, 2 monoecious genomes and 29 female genome alignments highlight contigs that are covered exclusively in male plants while having half of the coverage over other contigs in female genomes. These contigs with double coverage in females are believed to be X contigs while contigs with zero coverage in females are labeled as Y contigs (Figure 1).

To confirm this, Iso-Seq mRNA reads expressed in male flowers were mapped to the female reference. The ‘female-unmapped’ male mRNA reads were then mapped to the male reference to find male-specific mRNA expression with no homology to the female reference. These male-specific mRNAs were then intersected with the male-specific contigs to identify 574 genes on the non-recombining region of the Y chromosome. Prentout *et al.* has reported a similar approach using Illumina based RNAseq with Type I cannabis plants but the data is not currently available (Prentout 2019). It is important to note that reads that do not map to the female reference can be either 1) the Y chromosome and or 2) a structural variation (SVs) in the female reference genome. To avoid interpreting genome-specific SVs as Y associated, we considered only CNVs that exist in all males and females for X and Y categorization. Genes of interest on the Y chromosome include Enhanced Downey 2, FT Flowering Locus T, Flowering Time control protein FY, PIN2 (Auxin efflux carrier component 2), AP2-like ethylene-responsive transcription factor CRL5, and Protein trichome birefringence-like 6. These genes may play an important role in sex determinization, hermaphroditism, trichome development and day-neutral plant breeding and suggest careful selection of male genetics when culling males from breeding and propagation programs.

Previous assemblies from Laverty *et al*. and Grassa *et al.* focused on female Type I and Type III plants and used shorter reads delivering less contiguous assemblies and lower BUSCO scores (Grassa et al. 2018; Laverty et al. 2018). These studies leverage linkage maps to assemble contigs into scaffolds but this process does not rescue the missing BUSCO genes. It is important to note that deploying the same read lengths and quality on sibling genomes can present different results due to the variation in individual genome complexity and or the differing DNA isolation procedures used to extract the DNA. We chose not to use nuclei preparations in this study to ensure high coverage over the organellar genomes. One other male genome (Pineapple Banana Bubba Kush or PBBK) is deposited in NCBI with 18,355 contigs, 51Kb N50, 63% BUSCO completion and is smaller than the male genome presented here (512Mb vs. 876M-1Gb). The PBBK assembly is likely collapsed given its small size. The contigs of the PBBK assembly mostly map to female Jamaican Lion contigs. Of the unmapped contigs, only 60 Kb map to the Y chromosome. Given the low genome BUSCO completion numbers it is hard to know if this was truly a male genome or if the Y chromosome sequence has been collapsed with the X chromosome in the assembly.

### SNPs and indels across genomes

We used the DRAGEN unified genotyper to map and variant-call the 40 genomes against the maternal assembly (Miller et al. 2015). This produced 3.3 million to 19.3 million variants under 50bp in size (Figure 3) per cultivar. Mapping diploid whole genome shotgun data from a Jamaican Lion Mother Illumina® library results in 2.8 million SNPs where the most distant hemp sample (Merino) produces 17 million SNPs. This equates to a SNP every 312 bases in the diploid maternal reference and a SNP every 51 bases comparing Jamaican Lion to Marino Hemp. Other recreational Type I and Type II plants average 11.2 million SNPs and a sum of 12.8 million SNPs+Indels equating to a variant every 77-68 bases in most dispensary grade cannabis. To assess genotyping accuracy, Jamaican Lion Illumina® libraries were sequenced and mapped back to the Pacific Biosciences reference and 130,130 homozygous non-reference SNPs were detected suggesting a 4.2% false positive variant detection rate with short read mapping. As a comparison, Merino has 5.36 million homozygous SNPs detected. These variants were further annotated with SNPeff to identify 91,440 high impact male and female variants(Cingolani et al. 2012) (Supplemental Fig. S3).

**Figure 3:**
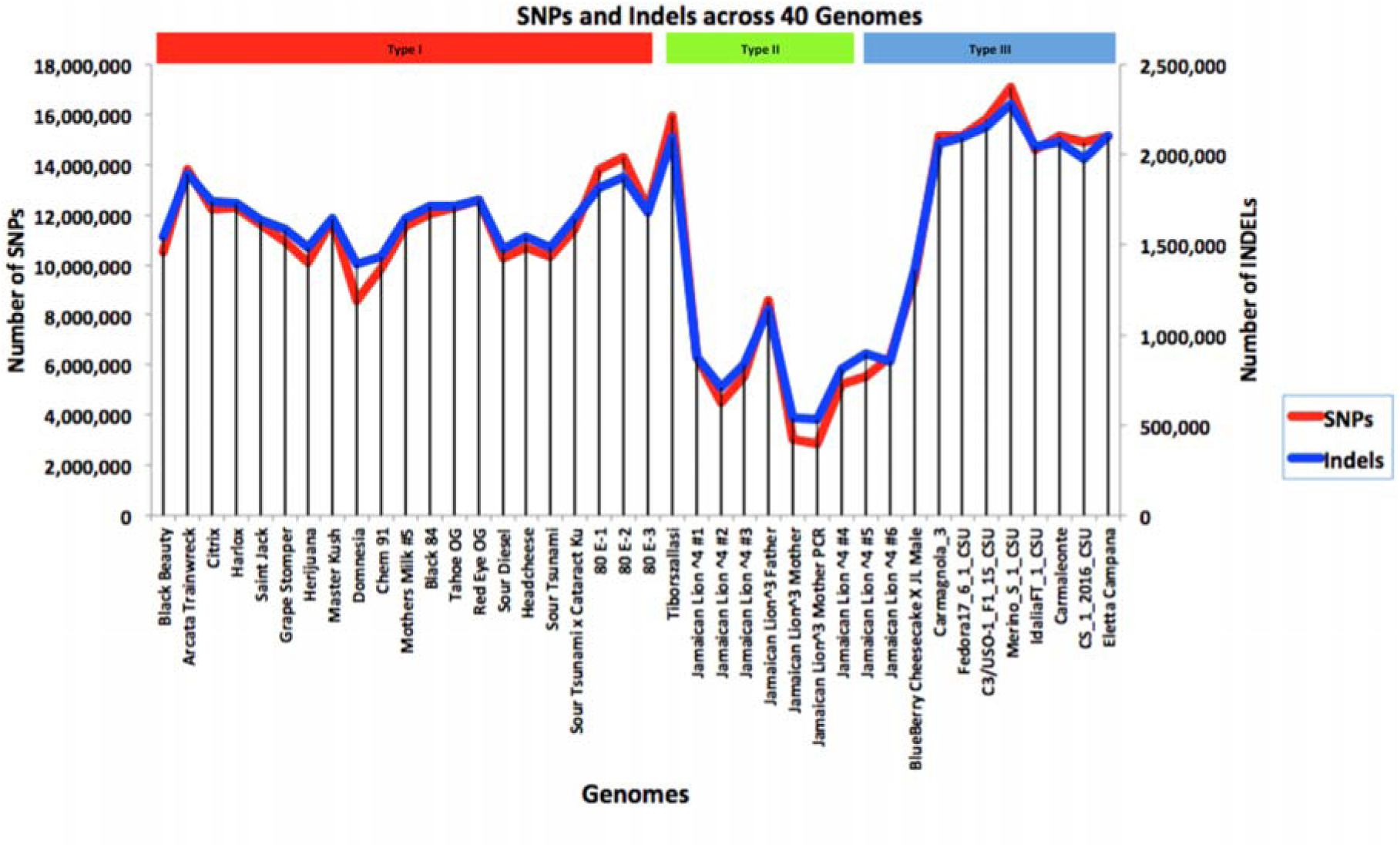
DRAGEN SNP calls across 40 genomes aligned to the maternal reference. DRAGEN conditions. 2 × 150 DNA sequences were aligned to the maternal Jamaican Lion reference using DRAGEN unified genotyping platform (DRAGEN HostSoftware V.05.011.281.3.2.5).

### Structural variation and gene models

Over 116Mb of structural variation were observed in the inbred trio with long reads using PBSV(https://github.com/PacificBiosciences/pbsv). In total, 27,664 female genes and 31,108 male genes were identified using 83,464 isoforms identified from the RNA derived from 5 male and female tissue types. Of these transcripts, 98.8% are observed in a recently published cannabis proteome from Orsburn *et al* (Orsburn 2019). Only 12,026 peptides were found in the mass spectrometry data suggesting low-level transcripts are likely below the sensitivity or sampling obtained with the platform. The distribution of structural variants in cannabis is non-uniform supporting the hypothesis of repeat driven genome plasticity described by Laverty *et al.* (Laverty et al. 2018) (Supplemental Fig. S4). In total, structural variations partially impact 1,446 genes in the trio.

### Copy number variation across the genomes

Copy number variation (CNV) in cannabinoid synthase genes has been reported previously (van Bakel et al. 2011; McKernan 2015; Weiblen et al. 2015; Pisupati et al. 2018; Vergara 2019). These studies were conducted with fragmented references or assays targeting specific genes. Whole genome analysis (52x average coverage) across highly contiguous references has not been completed to date. Illumina sequencing libraries were constructed using PCR (0 and 5 cycles for Jamaican Lion mother and 3 cycles for all other genomes) with unique molecular identifiers (UMIs) for deduplication of over-replicated molecules in the PCR process. The use of UMIs enables more robust copy number analysis with sequence data (Adams 2004). A PCR-free Jamaican Lion mother library was also constructed as a control (Jamaican Lion mother P). Coverage across 27,644 genes is 99.9% concordant between the PCR and PCR-free control libraries (Supplemental Fig. S5 & S6). The most discordant coverage was observed for JL5 (trio F1) and the three samples named 80E. These samples all exhibited extreme phenotypes (Figures 4 & 5). JL5 exhibited signs of dwarfism, short internodal spacing and stunted growth. The 80E samples have non-serrated leaf structures and powdery mildew resistance. Copy number variation is extensive in cannabis and is responsible for the chemical expression seen in Type I, II and III plants.

**Figure 4:**
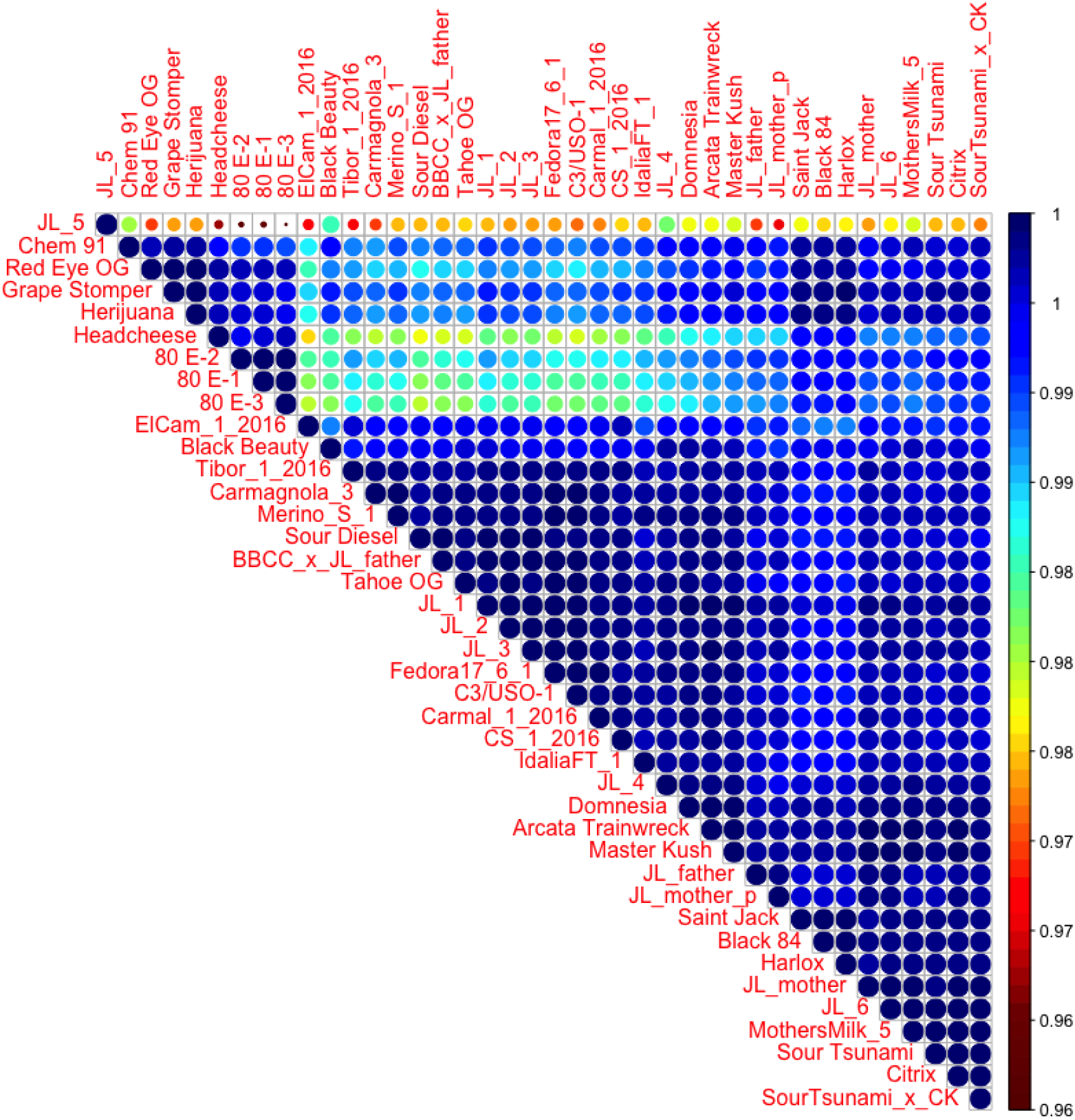
Coverage correlation matrix across 27,664 genes. Samples with lower coverage correlation (80e and JL5) have extreme phenotypes.

**Figure 5.**
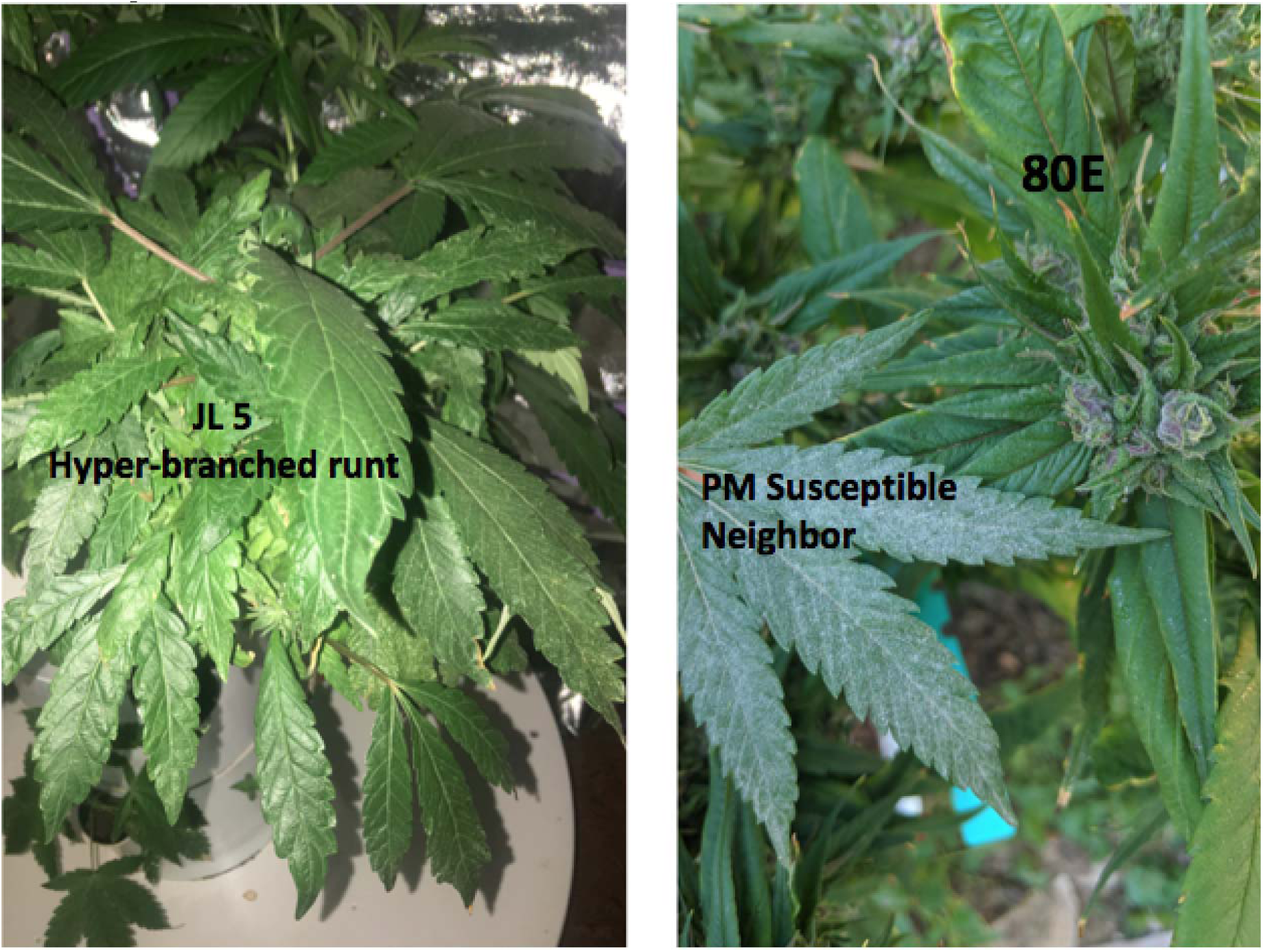
Samples JL5 (F1 Trio offspring sequenced with Pacific Biosciences) and 80E samples have the most discordant coverage variation and exhibit extreme phenotypes. JL5 (left) was termed a “runt” and exhibited signs of dwarfism, stunted growth, hyper-branching and short internodal spacing, despite being grown in the same hydroponic growth chamber as other siblings. 80E (right) samples were reported to be very resistant to powdery mildew and expressed unique non-serrated leaf structures. 80E can be seen on the right side of the right image with a neighboring powdery mildew infected plant.

### Pathogen response genes

*G. chicoracearum* has been shown to cause powdery mildew (PM) in cannabis while *Podosphaera macularis* has been reported to cause PM in *Humulus lupulus* L.(hop) which is a member of Cannabaceae and closely related to cannabis (Wolfenbarger et al. 2016; Punja 2018). Cannabis-derived powdery mildew can result in significant crop loss while exposing cannabis trimmers to powdery mildew-induced allergies (Victory et al. 2018; Zamir 2018). Many cannabis plants are believed to be powdery mildew-resistant but to date the genetics of this trait have not been described. Identification of the genetic basis of PM resistance can lead to more targeted breeding, increased yields and reduced employee allergen exposure. Cloning, expression and purification of the enzymes in a non-pathogenic bacterium could enable the development of foliar enzymatic sprays against epiphytic pathogens such as PM.

Copy number gains and losses in genes encoding three classes of resistance were evaluated. These data were compared to records from several cultivators on PM resistance of the submitted cannabis DNA samples. The existence of one or more copies of thaumatin-like protein (TLP) on contig 2563 was observed in several cannabis cultivars reported to be resistant to PM (Figures 6A and 6B). Endochitinase CH25 and lack of mildew resistance loci O (*MLO*) also correlated with resistance to PM (Supplemental Fig. S7). Several of these genes were heavily expressed in multiple tissues (Supplemental Fig. S9).

TLPs are responsible for a wide array of pathogen resistance in plants and have been reported to express PM resistance in *Vitis vinifera* (grape) and hops (Kappagantu et al. 2017a; Kappagantu et al. 2017b; Yan et al. 2017). TLPs copy number expansions in spruce are responsible for defense against *Botrytis* and other fungal pathogens (Šķipars 2017). TLP antifungal properties are believed to be due to their β-1,3-glucanase activity (Trudel et al. 1998; Grenier et al. 1999; Adams 2004). Genetic transformation of wheat with TLP and glucanases results in enhanced resistance to *Fusarium* (Mackintosh et al. 2007). Jongedijk *et al* (Jongedijk 2013) demonstrated synergistic activity of chitinases and β-1,3 glucanases in transgenic tomato. Given the complexity of the pathogen response in hops against PM, it is unlikely that a single gene is responsible for PM resistance in cannabis. Endochitinases, *MLO* and other pathogen response (PR) genes may augment or attenuate the response.

**Figure 6A:**
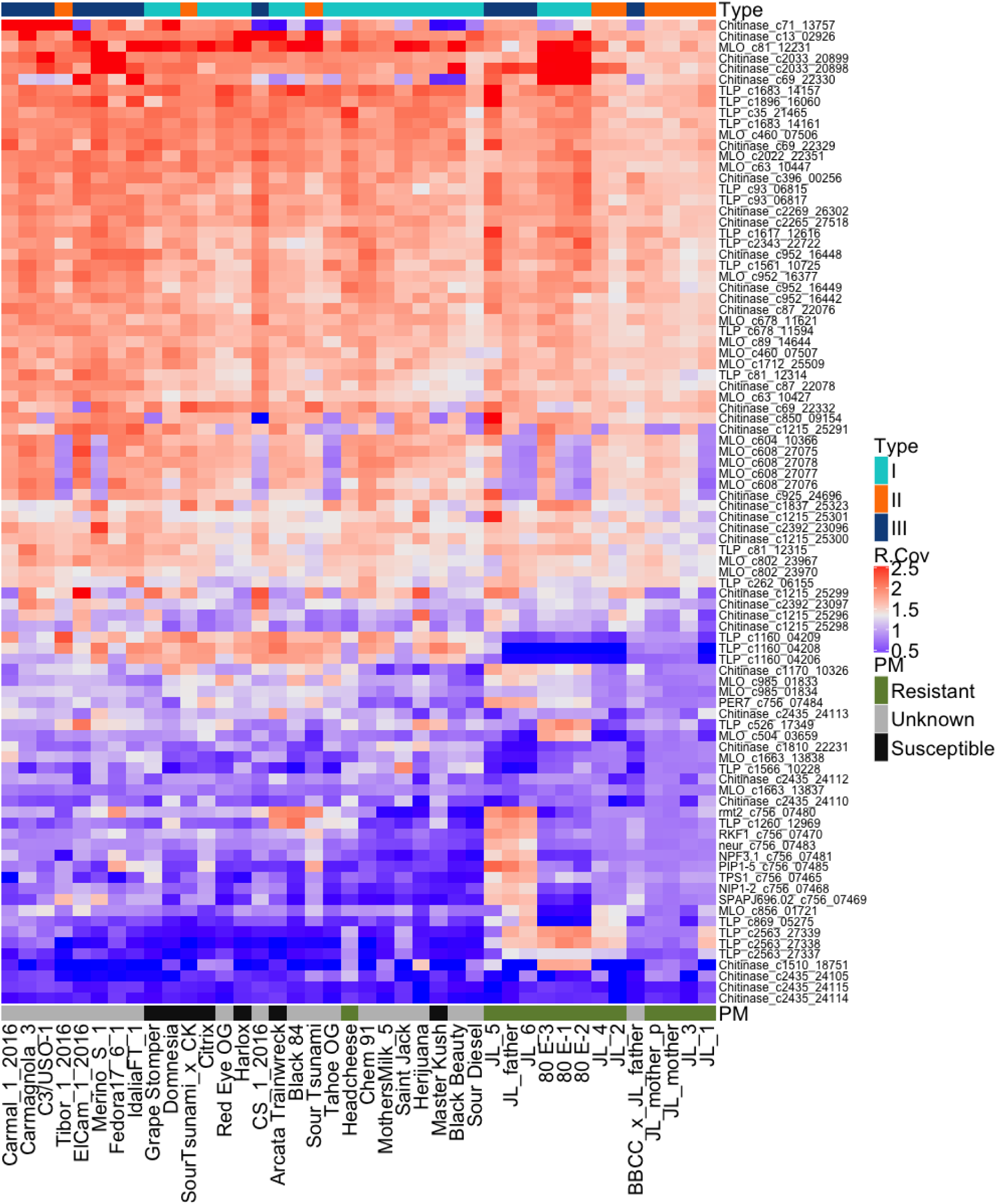
23 TLPs, 24 MLOs, 35 chitinase genes, pathogen response genes on the CBCA deletion, and their copy number variation across 40 cultivars. X-axis contains sample names and has two independent labels. The top label is Type I, II, and III classification. The bottom label is the reported powdery mildew-resistance status of each cultivar. *MLO* deletions likely increase resistance, while TLPs and chitinase deletions likely lower PM resistance.

**Figure 6B:**
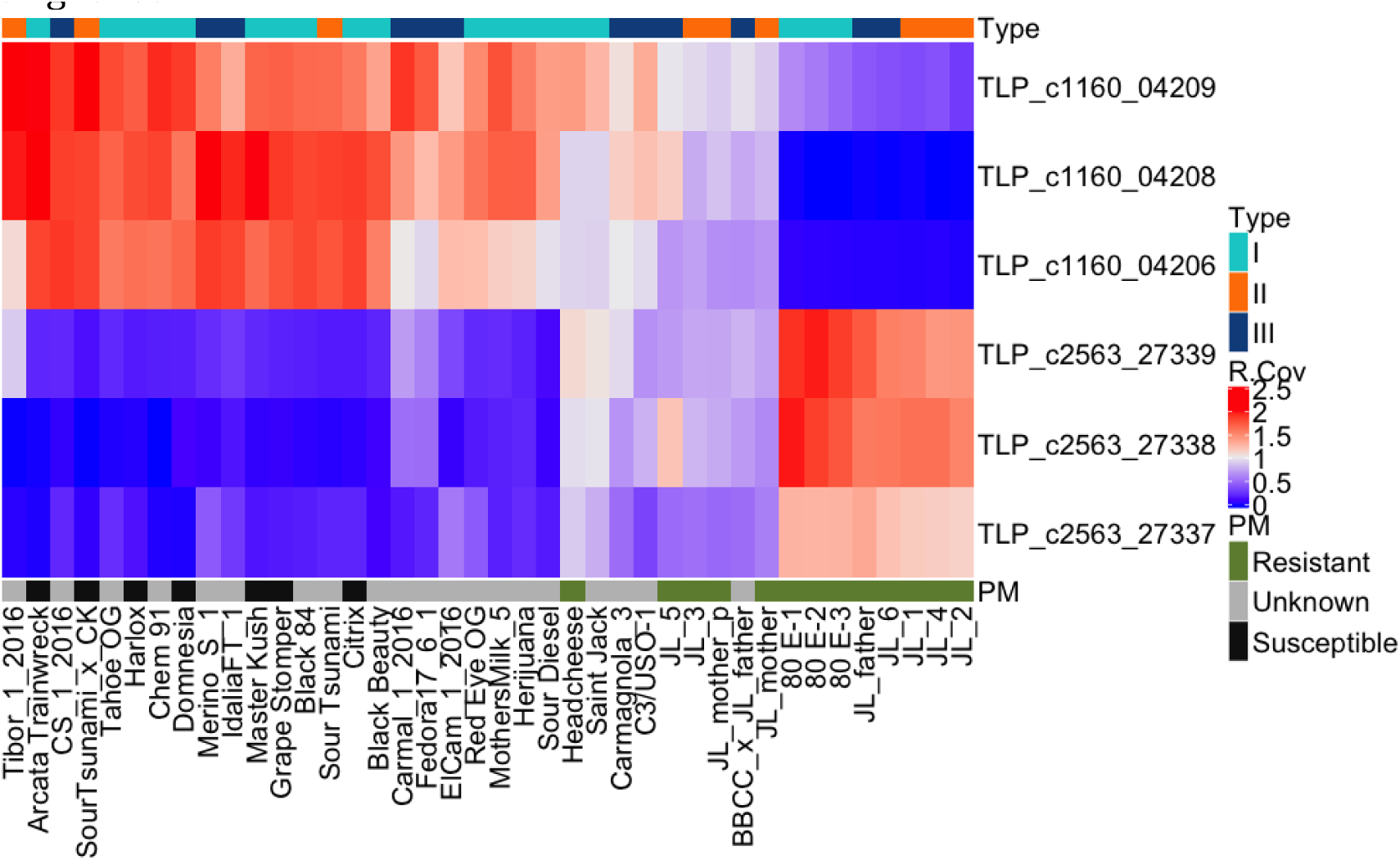
Copy number variation (CNV) of 6 TLPs normalized to whole genome coverage across 40 cultivars. X-axis contains sample names. The Y-axis contains 6 TLPs of interest. The top colored x-track contains Type I, II, II status of the samples while the bottom track contains the reported powdery mildew-resistance status. Powdery mildew-susceptible (S) cultivars cluster towards the left (Purple) and have a deletion of *Cs*TLP1. Unknown (U) powdery mildew-resistance status samples are in grey. Along the y-axis, 6 TLPs: deletions are blue in the heat map and amplifications are shown in bright red.

Twenty-three TLPs, 35 chitinases, and 24 *MLO* genes were found in the Jamaican Lion reference genome and were evaluated for gene expression in 5 parental tissues and genomic copy number variation across the 40 genomes. Many PM-susceptible cultivars reveal deletions of a TLP gene we have termed “*Cs*TLP1” while PM-resistant cultivars contain *Cs*TLP1 or copy number gains in *Cs*TLP1. RNA expression of *Cs*TLP1 was observed in all tissues except roots, with the highest expression in male flowers and female leaves. Due to the limited number of samples in the dataset and the presence of the Jamaican Lion family potentially producing synthetic associations, cloning and expression of putative resistance genes is necessary to confirm the role of *Cs*TLP1 (Goldstein 2011).

### Anti-fungal activity

*Cs*TLP1 was cloned into a pET-30a vector for expression in *Escherichia coli* for *in-vitro* fungicidal assays. Of the expressed and purified *Cs*TLP1 protein, 75% of the expressed protein was found in the inclusion bodies. *Cs*TLP1 was first confirmed to have β-1,3 glucanase activity using a malt β-glucanase assay (Megazyme). This was complemented with anti-fungal assays described by Misra *et al* (Misra et al. 2016) (Figure 7). Of interest is the visible reduction in red pigmentation of *Fusarium oxysporum* colonies. Red pigmentation in *F.oxysporum* has been reported to be the product of aurofusarin expression. Vujanovic *et al.* (Vujanovic 2017) describe a reduction in *Fusarium* aurofusarin expression with mycoparasitic and chemical control agents supporting the antifungal properties of *Cs*TLPs and chitinases.

*G.chicoracerum* (PM) is an obligate biotroph and is difficult to culture for controlled fungicidal evaluation of *Cs*TLP1. Instead, purified *Cs*TLP1 was applied to cultures of *Aspergillus flavus, Penicillium chrysogenum and Fusarium oxysporum*, which are other fungal pathogens of cannabis. Growth of *A. flavus* and *P chrysogenum* was not inhibited by *Cs*TLP1 but growth of *Fusarium oxysposum* was inhibited (Supplemental Fig. S10-12). Since TLPs often work synergistically with chitinases, co-application of *Cs*TLP1 and *T.viride* chitinase was explored. Co-application of these two enzymes inhibited both *P.chrysogenum* and *F.oxysporum* growth in vitro.

### Cannabinoid synthase genes

Thirty-nine cannabinoid synthase genes were evaluated for CNV across 40 genomes. Using the genomic DNA coverage maps across THCA synthase (THCAS), cannabidiolic acid synthase (CBDAS) and cannabichromenic acid synthase (CBCAS) (genes found on contigs 741, 1772, 756), we were able to classify plant primary cannabinoid expression into Type I, II, and III plants (Figure 9). These common deletions reflect the elusive Bt:Bd allele suggested by de Meijer *et al.*(de Meijer et al. 2003). Plants lacking a functional CBDAS gene are Type I plants. Type II plants have both functional genes and synthesize both THCA and CBDA. Plants with no functional THCAS gene and a functional CBDAS gene are Type III plants. Plants lacking both functional genes are Type IV plants and only synthesize the precursor cannabigerolic acid (CBGA). While deletions of entire THCAS and CBDAS genes are the most common Bt:Bd alleles observed, it is possible to have plants with these genes where functional expression of the enzyme is disrupted by deactivating point mutations (Kojoma et al. 2006). Further refinement of cannabinoid expression beyond the simple Type I –V designations may be augmented with RNA profiling.

**Figure 8.**
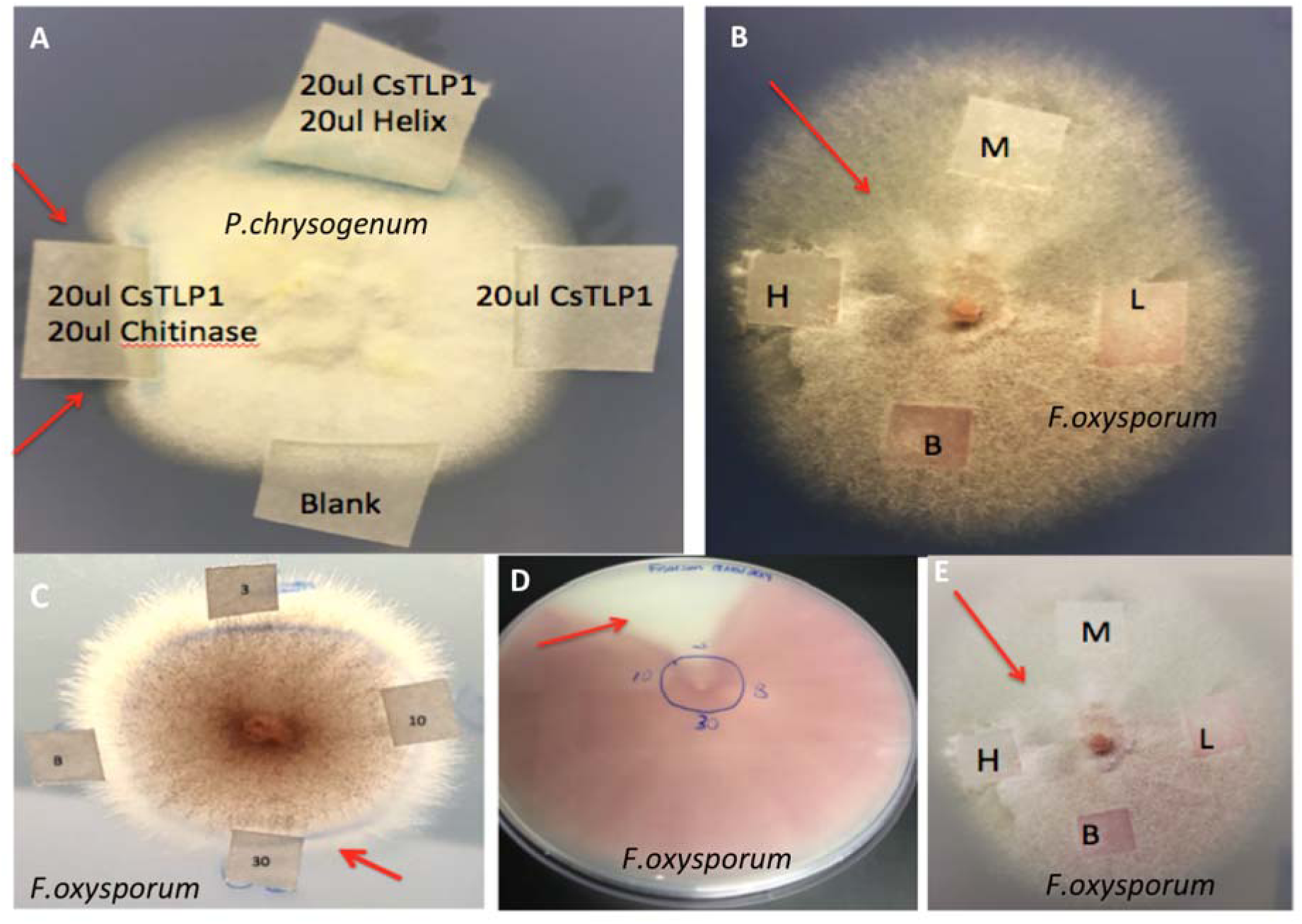
Antifungal activity of cloned *Cs*TLP1 using Misra et al. assay. A) *Penicllium chrysogenum* grown in the presence of dialyzed *Cs*TLP1, *Trichoderma viride* chitinase (Sigma) and a *Helix pomatia* β-1,3 glucanase. B) Enlarged image of *Fusarium* and *Cs*TLP1 (3 μg, 10 μg, 30 μg, and Blank) + *T. viride* chitinase (40 μL, 20 μL, 10 μL applied at 600 μg μL^-1^) plated on potato dextrose agar. C) *Cs*TLP applied only to *Fusarium* (3 μg, 10 μg, 30 μg, and Blank). D) Same colony as shown in lower left, two weeks later. In this case the 3 μg addition of *Cs*TLP demonstrated reduced pigment (aurofusarin) expression. E) Same image as B with different contrast to emphasize aurofusarin expression. Fungal cultures were incubated for three days to allow for development of a sizable colony and then assessed for disturbed growth (red arrows) after application of protein (*Cs*TLP or controls) on Whatman paper. Images were collected before and after the *Cs*TLP1 protein was applied and grown for another 36 hours. Subsequent to this, similar dosages of *T. viride* chitinase (3 μg, 10 μg, 30 μg, Blank; Sigma) were placed on the Whatman paper to identify synergistic effects.

**Figure 9.**
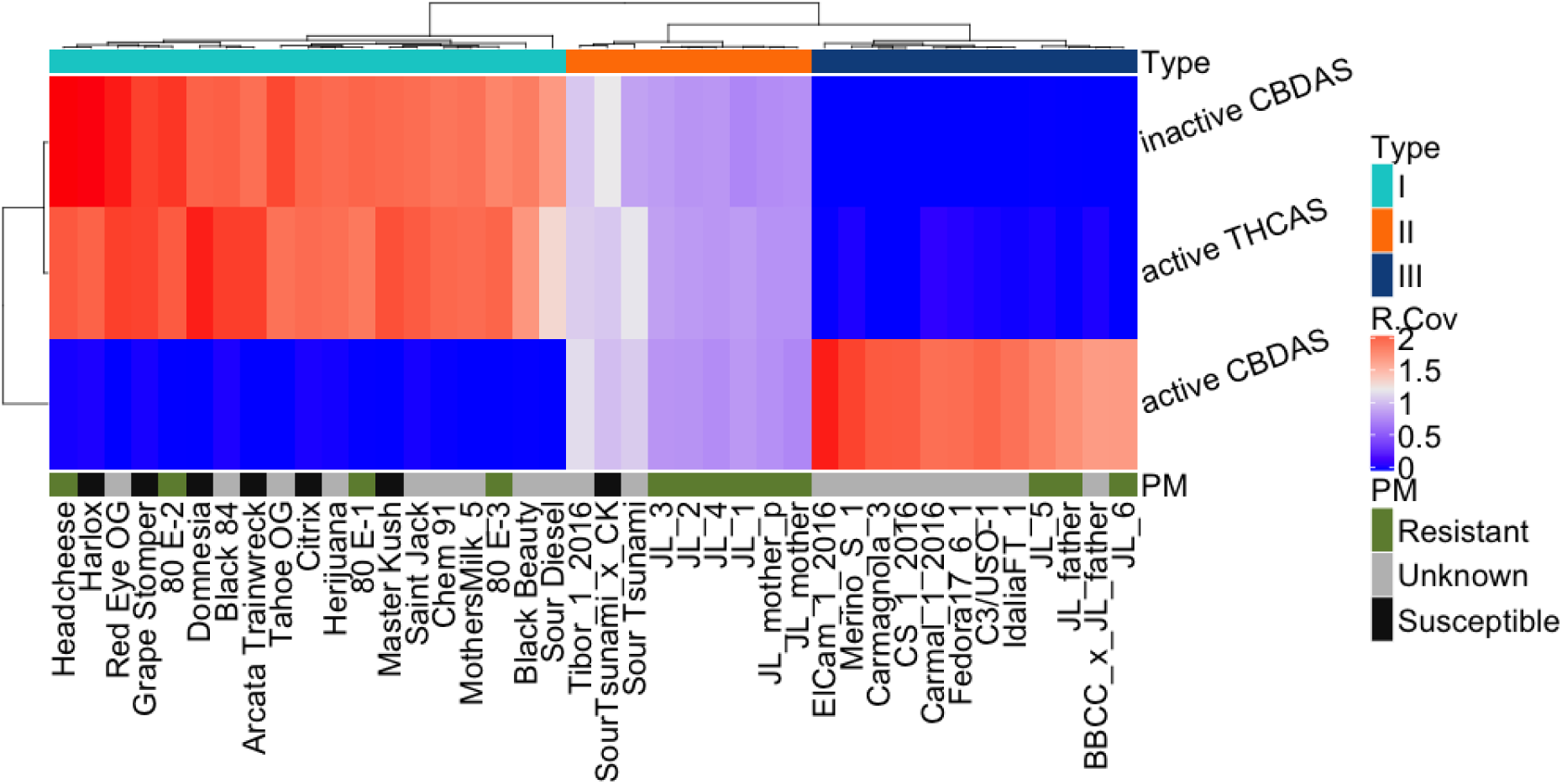
Copy number variation of the THCA and CBDA synthase gene normalized to whole genome coverage across 40 cultivars is shown. X-axis contains sample names. Y-axis contains the synthase gene names from contig742 and contig1772. Blue is deleted and red is diploid. Type I samples are on the right. Type II samples are in the center and Type III samples are on the left. The most common Bt:Bd allele across 40 genomes (x-axis) is a full gene deletion of THCAS or CBDAS. PM resistance genes are not in linkage with synthase genes.

Of interest is the frequent deletion of the CBCAS gene cassette (∼2Mb) seen on contig 756 (Figure 10). This contig contains 8 CBCAS genes directionally orientated and over 99.4% identical to each other. One CBCAS gene has recently been cloned and expressed and was previously known as “Inactive THCAS”(Laverty et al. 2019). Winnicki *et al.* demonstrated that multiple cannabinoids can be expressed from a single cloned synthase gene by modulating the yeast growth conditions (US patent 9,526,715 & 9,394,510)(Peet 2016) (Winnicki 2016). Hemp lines have also been more difficult to grow while maintaining a THCA concentration below the 0.3% THCA limit mandated in many jurisdictions. In particular, the THCA levels appear to increase in varieties from equatorial climates (Clarke 1996). Thus, it is possible that the presence of this cassette or other cannabinoid synthase CNVs are responsible for low levels of promiscuous THCA expression in some plants lacking a THCAS gene (Type III plants).

**Figure 10.**
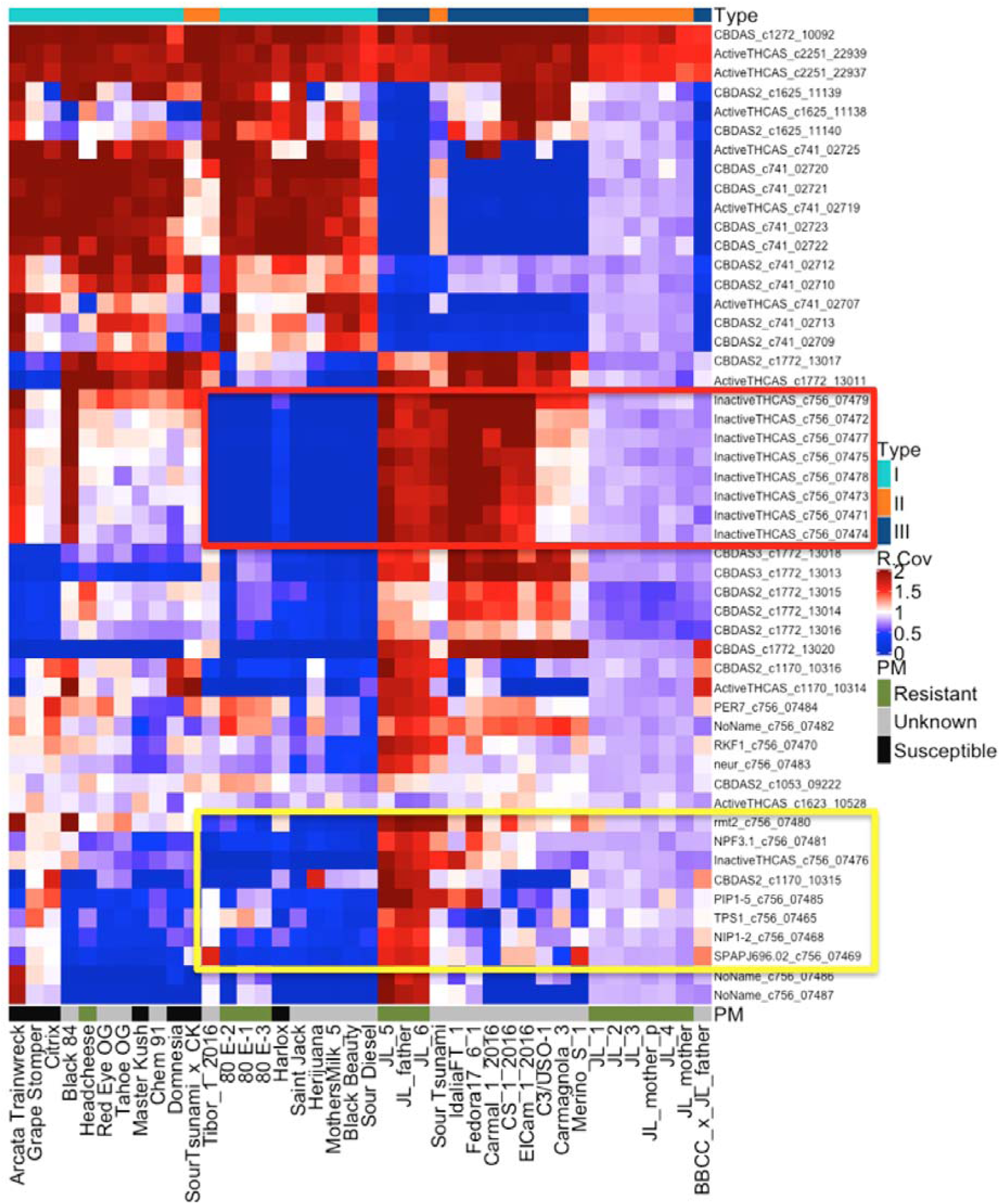
Copy number variation (CNV) heatmap of 40 genomes (x-axis) across 50 genes (39 cannabinoid synthase genes and 11 other genes including pathogen response genes) (y-axis). Note that many Type I plants lack a CBCAS or “Inactive THCAS” gene cluster (red box). Some Type I plants do exist with a CBCAS contig (Black 84 and Arcata Trainwreck) and most of the type III plants also have this contig (single exception is BBCC X JL father). Type II plants are usually heterozygous for this region. This large CBCAS deletion often contains pathogen response genes on contig 756 (yellow box) and segregates Type I plants via PCA analysis.

Additionally, the CBCA deletion also harbors pathogen response genes. The genes found in this deletion contain an expressed gibberellin transporter (NPF3) (Tal et al. 2016), RMT1 (involved in viral defense), PIP1 (PAMP-induced secreted peptides), and NIP1 (aquaporin involved in H_2_O_2_ pathogen response)(Carr and Loebenstein 2010; Sadhukhan et al. 2017). This implies that optimization of cannabinoid expression may need to be carefully monitored for pathogen susceptibility. Cannabinoid synthase CNV maps may play an important role in breeding for compliant pathogen-free hemp cultivars that do not synthesize residual THCA.

### Terpene synthase genes

Allen *et al.* made note of the very long terpene synthase (*Cs*TPS) introns in the maternal Jamaican Lion reference genomes (Allen et al. 2019). Longer genes are more prone to disruption by structural variations. Several *Cs*TPS genes have partial deletions captured in the PBSV VCF file. This led us to review copy number analysis over 22 *Cs*TPS genes (Supplemental Fig. S14). Significant copy number gains in *Cs*TPS17 are observed in multiple cultivars. Allen et al. classified *Cs*TPS17 as a potential myrcene or limonene synthase. Type II plants low in myrcene have been reported to be rare but in this analysis we observe the most extreme copy number variation in *Cs*TPS17 for Type II plants (Sour Tsunami, Tibor, JL1 –JL6 including the JL mother reference). Many *Cs*TPS genes have been reported to synthesize myrcene (*Cs*TPS3FN, *Cs*TPS30PK Booth et al. 2017) so the presence or absence of *Cs*TPS17 alone cannot predict complete chemotype. Nevertheless, the genome variation in *Cs*TPS genes does not appear to have undergone a bottleneck in Type II plants. Likewise, no *Cs*TPS genes are in close proximity to cannabinoid synthase genes, implying a weak linkage of CBDAS and THCAS with myrcene synthases. Type II plants being exclusively myrcene dominant is also challenged by a Limonene dominant Type II plant (Lemon Remedy) being in the marketplace since 2012.

### RNA Expression

Iso-Seq data was collected from parental tissues mainly to annotate the genome and capture high quality sequence of full-length transcripts for isoform annotation. While this data was incredibly valuable for genome annotation, its use for quantitation and comparative expression should involve more biological replicates. Additionally, the Iso-seq sequencing libraries do not appear to be saturated suggesting more sequencing with additional tissues and cultivars may discover more genes. The transcript counts for TLPs, chitinases, and *MLO*s are presented (Supplemental Fig. S9) demonstrating the highest transcript counts for *Cs*TLP1, chitinase_c2033, chitinase_c13, chitinase_c69 and chitinase_c87.

Some of the most extreme genome-wide and tissue-specific RNA expression is seen with chitinases. Of the differentially expressed tissues, chitinases are the top differentially expressed genes when comparing male flower RNA to female root RNA and female flower RNA compared to male flower RNA (CHIT5 on contig2033 or EFW9900020898). Chitinases were the second most differentially expressed genes when comparing female leaf RNA to female root RNA and when comparing female seeded flower RNA to male flower RNA (both CHIT5 or EFW9900020898). Chitinases were the third most differentially expressed gene (CHN14 on contig69 or EFW9900022332) when comparing female flower RNA to female seeded flower RNA. Low chitinase and TLP expression in roots may be required for commensal mycorrhizal interactions. The differential expression in female flower RNA to female seeded flower RNA was surprising. While these data are interesting they do not contain biological replicates. An improved whole genome enzymatic bisulfite method (EM-Seq) was explored in duplicate on 4 tissues to further understand transcriptional regulation.

### Methylation analysis

Four tissues were evaluated in duplicate with a novel enzymatic conversion assay utilizing APOBEC (Vaisvila *et al.* in press) known as EM-Seq (NEB). Like bisulfite sequencing, this method converts non-methylated cytosines into uracil but has more complete conversion, less GC bias in the library and generates longer insert size libraries. These features are important in cannabis as its genome becomes 83% AT after conversion and longer DNA molecules are easier to map to such low-complexity converted genome references. These data were compared to bisulfite treated control libraries combined with HPLC quantitation of digested nucleotides to quantitate conversion rates with both CpG methylated pUC and unmethylated Lambda DNA (Vaisvila *et al.* in press). We evaluated the CpG methylation 2kb upstream of the TSS for all chitinases and found Chitinase_c69 (EFW9900022332) was hypomethylated in both replicates supporting high transcription of this gene (Figure S15 & S16). Methylation data is also helpful differentiating NUPT (Nuclear Plastid) DNA from chloroplast DNA as the chloroplasts lack CpG methylation (Fojtova et al. 2001). Other notable differential methylation is observed between female seeded flowers and female flowers on the promoters of the Edestin gene involved in seed development and in THCAS between female flower tissues and male flowers (Docimo et al. 2014). THCAS expression is known to be highly expressed in female flowers and exhibit low expression in male flowers (data not shown).

### Yield related genes

Copy number analysis demonstrated JL5 and 80E samples have the most coverage discordance compared to all 40 cultivars. Further review of these copy number changes revealed a unique amplification of Gibberellic Acid Insensitive genes (GAI) (Supplemental Fig. S17) in the JL5 genome. These genes are reported to repress the gibberellin sensitive growth pathway and result in reduced stem elongation and dwarfism in Arabidopsis (Peng and Harberd 1993; Carol et al. 1995; Peng et al. 1997). Further cloning and validation is required to better understand these genes and their contribution to yield.

Using BCFtools we evaluated Mendel errors in the trio to be under 0.2% of the genotypes (1,763/1,173,954) (Narasimhan et al. 2016; Danecek and McCarthy 2017). While the SNP analysis of the Jamaican Lion strains supports a high relatedness between JL5 and its parents, somatic variation in flowering plants is known to be age related and exacerbated with transposon rich sequences (Singh et al. 2015). Just upstream of GAI is a Gypsy LTR that may be responsible for somatic variation in this offspring. Further work is required to better understand the mutagenic potential of cannabis embryogenesis.

## Methods

### DNA Purification

High molecular weight DNA was extracted from isopropanol-treated stalk and leaf, utilizing a modified CTAB, chloroform and SPRI technique. Plant material was frozen at −80°C and ground to a fine powder with a mortar and pestle. Next, 100mg of plant material was aliquoted into two mL tubes with one mL MIP Solution A (Medicinal Genomics, Beverly MA) and placed on a rotator for 30 minutes at room temperature. Then, 4 µL RNase-A (4 mg-mL^-1^, Promega, Madison WI) was incubated at 37 °C for one hour vortexing every 15 minutes. Subsequently, 5 µL Proteinase-K (20 mg ml^-1^, Qiagen) was added to the solution and incubated at 60°C for 30 minutes, vortexing every 10 minutes with a 10 minute incubation on ice in between vortexing events. After centrifugation for 5 minutes at 14,000 rpm, 600 µl of the supernatant was removed and added to 600 µL chloroform. This solution was vortexed until it turned milky white and was then spun at 14,000 rpm for five minutes. Four hundred µl of the aqueous layer was removed with a wide bore tip and put into a 1.5 ml tube with 400 µl of MIP solution B (Medicinal Genomics). This solution was gently inverted and 400 µl of MGC magnetic binding buffer (Medicinal Genomics) was added and the tubes were placed on a room temperature rotator for 15 minutes. The tubes were then placed on a magnet stand for three minutes and washed three times with one ml of 70% ethanol. Beads were dried for five minutes at room temperature and eluted at 56°C in 100 µl of 10 mM Tris-HCl. The eluted samples were then incubated at 56°C for five minutes, returned to the magnetic rack, and the eluted DNA was removed from the binding beads. Samples prepared in multiple tubes were pooled and evaluated on a Qubit®, Nanodrop® and an Agilent gDNA Tape Station®.

Twenty µg of high molecular weight (HMW) DNA (50 ng µl^-1^) was evaluated on a Fempto Pulse (Agilent) with a three hour run time and converted to SMRTbell® libraries using SMRTbell® Express Template Prep kit according to the manufacturer instructions. In total, 91 Gb of data was generated across 11 SMRT cells with Sequel SW 5.1 and Polymerase binding/ Sequel Sequencing Kit 2. Three more SMRT cells were loaded with Sequel SW 6.0 and polymerase Binding 3.0/Sequel Sequencing Kit 3.0. Twenty-hour movies were recorded generating an additional 34 Gb of sequence (Table 1B).

**Table 1A.** Pacific Biosciences coverage and sequencing statistics of three Jamaican Lion cannabis genomes. Genomes were sequenced with continuous long read mode (CLR). F1 was female. BUSCO: benchmarking universal single-copy orthologs. Software and database versions: BUSCO.py 3.0.2, Augustus 3.3.21, Hmmer 3.2.1, Blast 2.7.1, eudicotyledons_odb10(Simao et al. 2015).

| **Coverage** | **Mother** | **Father** | **F1 (JL5)** |
| --- | --- | --- | --- |
| N50 | 3,283,100 | 1,668,042 | 3,491,975 |
| Contig number (> 5 Kb) | 481 | 1264 | 658 |
| Genome size (bp) | 875,793,298 | 1,009,156,132 | 999,122,115 |
| Complete BUSCOs (%) | 96.1 | 97.0 | 97.3 |
| Single-copy BUSCOs (%) | 83.5 | 63.3 | 63.5 |
| Duplicated BUSCOs (%) | 12.6 | 33.7 | 33.8 |
| **Sequencing statistics** |  |  |  |
| Unique molecular yield (Gb) | 125 | 150 | 84.8 |
| N50 RL (Kb) | 34.6 | 35.6 | 50 |
| N50 Subread (Kb) | 20 | 24 | 19 |

**Table 1B.**
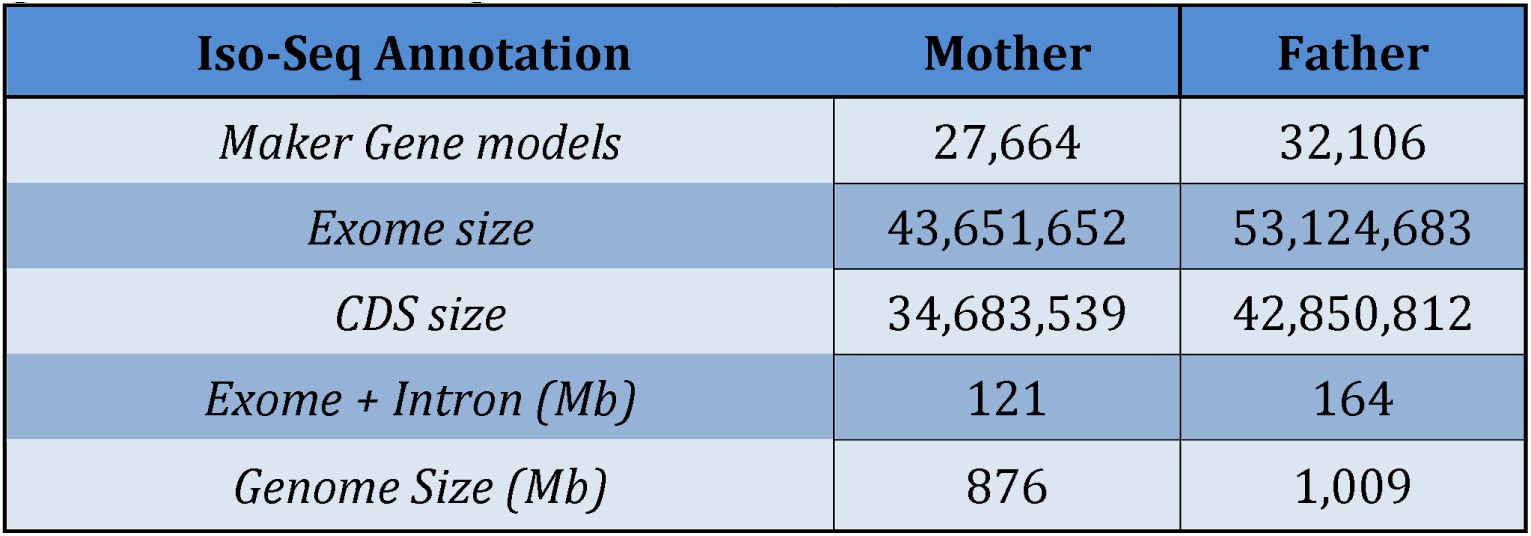
Estimated Exome size of Male (Father) and Female (Mother) Jamaican Lion genomes. The Father genome has an additional 118Mb Y chromosome.

### RNA purification

500mg of tissue was carefully diced into 40ml of 4°C RNALater (Sigma) for transport and storage. RNA was purified using a Monarch RNA purification kit (NEB). RNA quality was evaluated with an Agilent Tape station. Over 100ng/ul of RNA was purified with RIN numbers from 5.5-8.1. 1ug of purified poly A selected RNA was utilized to generate IsoSeq (Pacific Biosciences) according to the manufacturers instructions.

### Sequencing and mapping of 40 genomes

Illumina® whole genome sequencing libraries were prepared with the NEBNext Ultra II FS kit according to the manufacturer protocol. One hundred ng of cannabis genomic DNA was treated with the fragmentation enzyme mix for eight minutes at 37 °C. NEBNext® Multiplex Oligos for Illumina®(Unique Dual Index UMI Adaptors DNA Set 1) were ligated to the fragmented DNA by Ultra II ligation master mix at 20 °C for 15 minutes. The adaptor-ligated DNA was then size selected and PCR amplified using three cycles (all genomes except for Jamaican Lion mother) and five cycles for the Jamaican Lion mother with NEBNext® Q5 master mix. The final products were further purified with 0.8× NEBNext Sample Preparation Beads. Libraries were quantitated and size-evaluated with TapeStations®, normalized, and pooled. The pools were sequenced on the Illumina® Novaseq S4 using the XP workflow for paired-end reads of 2 × 150 cycles. Additionally, we constructed and sequenced a PCR-free Jamaican Lion mother control library using these same methods.

Two × 150 DNA sequence was aligned to the maternal Jamaican Lion reference using DRAGEN unified genotyping platform (DRAGEN HostSoftware Version 05.011.281.3.2.5).

Commands used for mapping and genotyping were “dragen –r –enable-variant-caller true –vc –vcmit-ref-confidence GVCF –duplicate-marking true -–enable-map-align-output true” and “dragen –f –r –enable-joint-genotyping true”.

### Pacific Biosciences Sequencing of the Trio

SMRTbell® sequencing libraries were constructed by the service provider according to the manufacturers instructions and produced 84.8 Gb to 125 Gb of unique molecular yield for Mother, Father, F1 genomes (Table 1B). These were assembled into 1.6Mb to 3.8Mb N50 contigs with the FALCON Unzip assembler v.0.3.0 (Table 1A).

### DNA Assembly

Falcon Unzip was configured as follows:

input_type = raw

genome_size = 1000000000

length_cutoff = 35000

length_cutoff_pr = 30000

pa_daligner_option = -e0.76 -l5000 -k18 -h480 -w8 -s100

ovlp_daligner_option = -k24 -h480 -e.95 -l5000 -s100

pa_HPCdaligner_option = -v -B128 -M24

ovlp_HPCdaligner_option = -v -B128 -M24

pa_HPCTANmask_option = -k18 -h480 -w8 -e.8 -s100

pa_HPCREPmask_option = -k18 -h480 -w8 -e.8 -s100

pa_REPmask_code=2,10;0,300;0,300

pa_DBsplit_option = -x500 -s400

ovlp_DBsplit_option = -s400

falcon_sense_option = --output_multi --min_idt 0.70 --min_cov 4 --max_n_read 200 --n_core 24

overlap_filtering_setting = --max_diff 100 --max_cov 250 --min_cov 4 --n_core 24

A single round of Arrow was used to polish the assemblies (Chin et al. 2016). Quast 5.0 was used to calculate assembly statistics (Gurevich et al. 2013; Mikheenko et al. 2018). BUSCO was utilized to measure a 96.1%, 97.0% and 97.3% complete ortholog assembly for the father, mother, and F1 genome respectively (eudicotyledons_odb10 lineage: Table 1A)(Simao et al. 2015; Waterhouse et al. 2017). Several attempts at utilizing Purge Haplotigs were made with varying minimap stringency scores (data not shown). These purged assemblies also purged the CBCAS synthase gene cluster, so we opted to only purge the maternal reference with the less stringent default settings. Cytogenetic estimates of cannabis genome size range from 0.84 pg to 0.91 pg suggesting a genome size of 821 Mb to 919 Mb (Divashuk et al. 2014). Running Polar star and Purge Haplotigs on the 3.8 Mb maternal reference resulted in a haploid assembly size of 876 Mb with a 3.2Mb N50.

### Gene annotation

Repeats were first classified by RepeatModeler. This identified 1,195 repeat sequences (37 bp to 16,027 bp. average = 1,381 bp, median=591 bp). Three rounds of MAKER2 were deployed. The first round utilized Iso-Seq mRNA data matched against four Rosaceae protein sets (European pear, wild strawberry, China rose, and apple). *Ab initio* gene prediction was trained with SNAP (hidden Markov-models or HMM-based gene finder). The second round of MAKER2 utilized the repeat database combined with EST alignments and BUSCO/Augustus cannabis-specific HMMs. This output was used to retrain SNAP. The third round of MAKER2 filtered the Annotation Edit Distance (AED = 0 to 1). SNAP was re-trained on this model and produced approximately 60,000 gene models where 24,000 of which had Iso-Seq support. Filtering this set on AED<1 with Interproscan support delivered 27,644 female gene models and 32,108 male gene models.

### Copy number analysis

Copy number analysis was conducted with BEDtools and the R statistical programming language(Quinlan 2014). Briefly, the per-base depth across the Jamaican Lion maternal reference genome was calculated using BEDtools genomecov command on mapped Illumina reads. Per-base depth for 27,664 genes was extracted using BEDtools intersect command with the Jamaican Lion maternal reference gene annotation file. Coverage of each gene was calculated by averaging the depth of each gene, and then normalized with mean depth across the genome for each sample. The normalized mean coverage for each gene of each sample was processed and compared using R packages including heatmap2, reshape2, dplyr, gplots, matrixStats, and factorexta.

### EM-seq libraries and Methylation analysis

Fifty nanograms of cannabis genomic DNA (with spike-in controls: unmethylated lambda DNA and CpG methylated pUC19 DNA) was sheared to 300bp with a Covaris S2. This DNA was end repaired and adaptor ligated utilizing the NEBNext EM-Seq protocol according to the manufacturers instruction. Six PCR cycles were utilized to amplify the libraries. Libraries were constructed in duplicate. One hundred million 2 × 76 bp Illumina sequencing reads were collected per sample on a Illumina NextSeq. Reads were mapped using the DRAGEN methylation mapper version 3.4.5 (Illumina).

### Chromosome structure

Two other cannabis assemblies have been published with suggested chromosome structures (Grassa et al. 2018; Laverty et al. 2018). Prentout et al. (2019) has highlighted that chromosome 1 in Laverty et al. is actually the X chromosome while SynMap2 alignments of Jamaican Lion to CBDrx have CBDrx chromosome six as the third largest chromosome in Jamaican Lion (Supplemental Fig. S2). In an effort to identify the source of these differences (biological vs. assembly artifacts), over 600 million HiC reads were generated with Phase Genomics using a modified DNA purification process (no nuclei preparation). This modified library protocol produced lower than average long-range links. Both SALSA and Promixo were used to estimate scaffold order and orientation of contigs (data not shown) but these data had a low concordance with Oxford Nanopore scaffolding (ONT). Hi-C data (generated on Jamaican Lion) and low coverage linkage mapping data (seen in Laverty and Grassa) utilized short read Illumina data. These shorter reads suffer read mappability issues in regions with tandem LTR sequences. ONT data has lower quality in the context of inverted repeats (Spealman 2019), but many of the cannabinoid synthase genes are separated by tandem LTRs. The cytogenetic data on cannabis chromosomes demonstrates the chromosomes have very similar sizes (Divashuk et al. 2014). Given the high frequency of 2Mb deletions and the very large size of the structural variations between cultivars discovered by PBSV, it may be premature to draw a consensus chromosome nomenclature for the entire cannabis species until more reference grade genomes are surveyed. We have opted to leave this reference un-scaffolded until more convincing data is generated on the Jamaican Lion chromosome structure. It should be re-iterated that each genome project targeted a different cannabis variety (Type I with Laverty, Type II with Jamaican Lion and Type III with Grassa) thus lack of consensus on the topic may be reflective of real biological variation.

### *Cs*TLP expression

A 225 amino acid (MW = 23,714.5) peptide (*Cs*TLP) was expressed in *E. coli* DH10B in 1L of terrific broth (TB, 24 g L^-1^ yeast extract; 20 g L^-1^ tryptone; 4 mL L^-1^ glycerol; 0.017 M KH_2_PO_4_, 0.072 M K_2_HPO_4_) with a 6× Histag. A 20 amino acid N-terminus signal peptide was removed. The amino acid sequence was: MIQNNCGRTIWPATQSGSGSSQLSTTGFELASGASQSIEIPAGPWSGRFWGRDGCSTDSSGRFA

CASGDCASGTVECNGAGGVPPTTLVEITVAENGGQDFYDVSNVDGFNLPVSVRPEGGNGDCQESTCPNNLNDGCPADLQYKSGDDVVGCLSSCAKYNMDQDCCRGAYDSPDTCTPSESANYFEQQCPQAYSYAYDDKTSTFTCSGGPNYLITFCPHHHHHH.

*Cs*TLP1 was eluted in 20 Mm Tris, 500 mM NaCl, 10 mM reduced glutathione (GSH), 1 mM glutathione disulfide (GSSG), 20 % glycerol, pH 7.5.

### β-1,3-glucanase assay

A malt β-D-glucanase assay (Megazyme) was used to measure *Cs*TLP1 enzyme activity according to the manufacturer protocol. The protocol was scaled down to fit in a 1.5 mL Eppendorf tubes. One hundred and fifty µL of dye-labeled azo-barley glucan was mixed with 150 µL of enzyme and incubated at 30 °C for 10 minutes. Nine hundred µL of precipitation solution A was used to precipitate undigested glucan. Samples were centrifuged for 10 minutes at 1000 × g to pellet digested glucan. Dye-labeled digested glucan remained in the supernatant and absorbance was measured at 590 nm.

### Chemotyping

Fourteen cannabis samples acquired within the state of Massachussets had HPLC measurements for CBDA/CBD and THCA/THC on the label. All chemotypes agreed with the genetic designations obtained with the Bt:Bd allele deletion status. While the deletion status of CBDAS and THCAS can predict general CBD:THC ratios as described by McKernan *et al.*(McKernan 2015), it cannot segregate 0.3% THCA from 0.5%THCA-producing Type III plants. It is hypothesized that residual THCA production in Type III plants that have no THCAS gene are the result of promiscuous expression of THCA from other cannabinoid synthase genes. CBDAS and CBCAS have the highest sequence and amino acid similarity to THCAS.

## Conclusions

Cannabis exhibits complex variation. Millions of SNPs and indels and thousands of CNVs and other structural variations are important data to evaluate for effects on commercially meaningful chemotype expression. This variation can be seen as both a blessing and a curse. While diversity is expected given the complex chemotypes seen in cannabis, stable seed lines will be more difficult to produce without a better understanding of apomixis potential in cannabis. Likewise the natural diversity observed and complicated copy number amplifications of various key chemotypic genes may favor accelerated breeding approaches over multi-target or multi-megabase CRISPR editing and transformation.

These data also address a long-standing question regarding monoecious chromosome structures assessed by Razumova *et al.* (Razumova et al. 2016). Monoecious varietals (C3 and Fedora) cluster together in Figure 1 and the whole genome shotgun data clearly demonstrates an XX chromosome structure for monocieous samples thus further verifying earlier cytogenetic assessments. This has been challenging to conclusively determine with genetic markers (MADC2 and SCAR) developed prior to any knowledge of a genomic reference. The pioneering primers described by Mandolino et al. are now known to have homology to many autosomal repeat structures in the genome (Mandolino 1999) and may lead to false positive or false negative results when applied across cannabis diversity. Primers with higher male specificity are more easily assessed with multiple genomes comprehensively sequenced.

These data also resolve the equally evasive Bt:Bd allele suggested by de Meijer et al(2003). While much work has been published on these hypothetical alleles, their precise genomic coordinates could not have been resolved without long read sequencers capable of spanning the complicated tandem repeat structures observed in the cannabinoid synthase gene clusters.

Of particular relevance to recent hemp and cannabis legislation are the insights regarding why Type III hemp plants lacking a THCA synthase gene can still synthesize small amounts of THCA. Type III plants, lacking a functional THCAS gene but containing a functional CBDAS gene, frequently have a CBCAS gene cluster. This CBCAS gene cluster is not seen in all Type I plants (plants lacking a CBDAS gene but containing a THCAS gene). This CBCAS gene cluster shares the most sequence similarity with THCAS and CBDAS and may produce THCA as a byproduct. Breeding for plants that contain the natural CBCAS deletion in the population will also select for plants lacking the linked pathogen-resistant genes (NIP1, PIP1, NPF3, RMT1). Breeding for less than 0.3% THCA production may enrich for pathogen susceptibility and higher patient fungal exposure.

While yeast models support byproduct production by synthase genes, more work is required to confirm THCA production from CBCAS or other cannabinoid synthase genes *in planta*. This is important to consider given the documented but rare cases of cannabis acquired aspergillosis fatalities while human exclusive THCA toxicity has never been conclusively documented (McKernan et al. 2015; McKernan et al. 2016; McKernan 2018b). While, the recent vaporizer fatalities are associated with THCA products, these toxicities are not attributed to the therapeutic index of THC but to the solvents (Vitamin E acetate) often used in the clandestine manufacturing practices(Blount et al. 2019).

This work represents preliminary functional validation of pathogen resistance genes in cannabis. Compensatory pathogen response pathways like the 24 MLO loci and 35 endochitinase genes need further cloning and scrutiny. Co-transformation of chitinases and TLPs have shown increased pathogen resistance to *Sclerotinia sclerotiorum* in *Brassica* (Aghazadeh 2016). TLPs, chitinases and lack of certain *MLO* genes are preliminary markers for pathogen resistance and may accelerate breeding for more resistant lines.

There are several limitations to this study. First, powdery mildew susceptibility is not a binary trait and some plants may only show susceptibility under stress. Second, several different modes of resistance are possible. While further functional studies are required to establish definitive PM resistance markers, highly expressed TLPs and Chitinases are promising targets. This refined genetic map of the variation in cannabis can guide more stable and directed breeding efforts for desired chemotypes and pathogen resistant cultivars. A refined map of the Y chromosome may also lead to a better understanding of the inheritance of hermaphroditism and possibly the production of double haploid seeds.

A comprehensive understanding of the diversity of cannabinoid synthesis genetics may assist in resurrecting rare cannabinoids bred out of the population during prohibition. Most jurisdictions incarcerated based on the weight of cannabis plant matter seized and did not take note of the potency of the product. Much like during the prohibition of alcohol, beer production was replaced with higher concentration whiskeys and spirits. Similarly, cannabis prohibition selected for the production of more THCA per kilogram of product. This incentivized the selection of unseeded flowers from Type I cannabis plants. This selection is believed to have come at the cost of Type II and Type III genetic diversity. These data suggest analogous selections for CBCAS deleted cultivars may occur with the 0.3% THCA hemp limits being imposed in many jurisdictions and this selection may result in plants with deleted pathogen response genes. Future studies may entail measuring the pathogen burden of type III plants with and without the CBCAS deletion to better understand the tradeoffs these THCA limits impose.

With the diversity of cannabinoid synthesis genes being rapidly discovered combined with novel phytocannabinoids being discovered every year, the Type I – Type V cannabis nomenclature system will likely require further refinement (Citti 2019). We believe these data may inform the process.

## Data availability

Data is available at NCBI under Project ID PRJNA575581 and SUB6635057. A genome browser is available at CoGe (https://genomevolution.org/) A phylogenetic blockchain registration tool is available at Kannapedia.net. This platform hosts public access to an additional 420 samples sequenced with a 3Mb Agilent SureSelect panel that includes the Bt:Bd allele.

## Author contributions

KJM-Designed study, grant application, field DNA isolation, cultivation advice, data analysis, anti-fungal assays, cryptocurrency methods, manuscript writing YH-DNA and RNA purification, Library construction, anti-fungal assays, DNA sequencing

LTK-DNA and RNA purification, Library construction, DNA sequencing

HE-DNA purification

LZ-Library construction, DNA sequencing

ZE-Cultivation advisor

MJ-80E breeding and tissue culture

SK-FALCON-Unzip Assembly

PB-DNA Sequencing strategy and DNA purification advice

MP-Proximo Assembly Hi-C advice

WB-DNA shotgun and assembly advisor

TH-DNA shotgun and assembly advisor

## Acknowledgements

Luo Sun, Eileen T. Dimalanta, Brittany S. Sexton, Keerthana Krishnan, Bradley W. Langhorst of New England Biolabs for methylation, sequencing, and library construction advice.

Jeremy Menard for his breeding expertise Maximillian Press for Proximo Assembly Hi-C advice Colorado Seeds for Powdery Mildew resistance expertise

Seth Crawford for experience with CBCA deletions and Type IV plant breeding

Zamir Punja for cannabis pathogen expertise

NETA for Powdery Mildew resistance expertise

Champlain Valley for Powdery Mildew resistance expertise Polaris Wellness for Powdery Mildew resistance expertise John MacKay for hemp genetics and DNA purification.

Parabrick for GPU variant calling comparisons to DRAGEN. R.T.M for his brave fight with cancer and advice on this matter. Rebecca McKernan for editorial advice.

Kyle Boyar for grant review. Rachel Backer for careful editing

This project was partially funded by a blockchain grant from the Dash DAO.

Solarguy, Mastermined, Mark Mason, Joel Valenzuela, TanteStefana, Coingun and the

Dash masternode network for their advice and guidance on the Dash proposal process.

## Supplemental Data

**Supplemental Figure S1:**
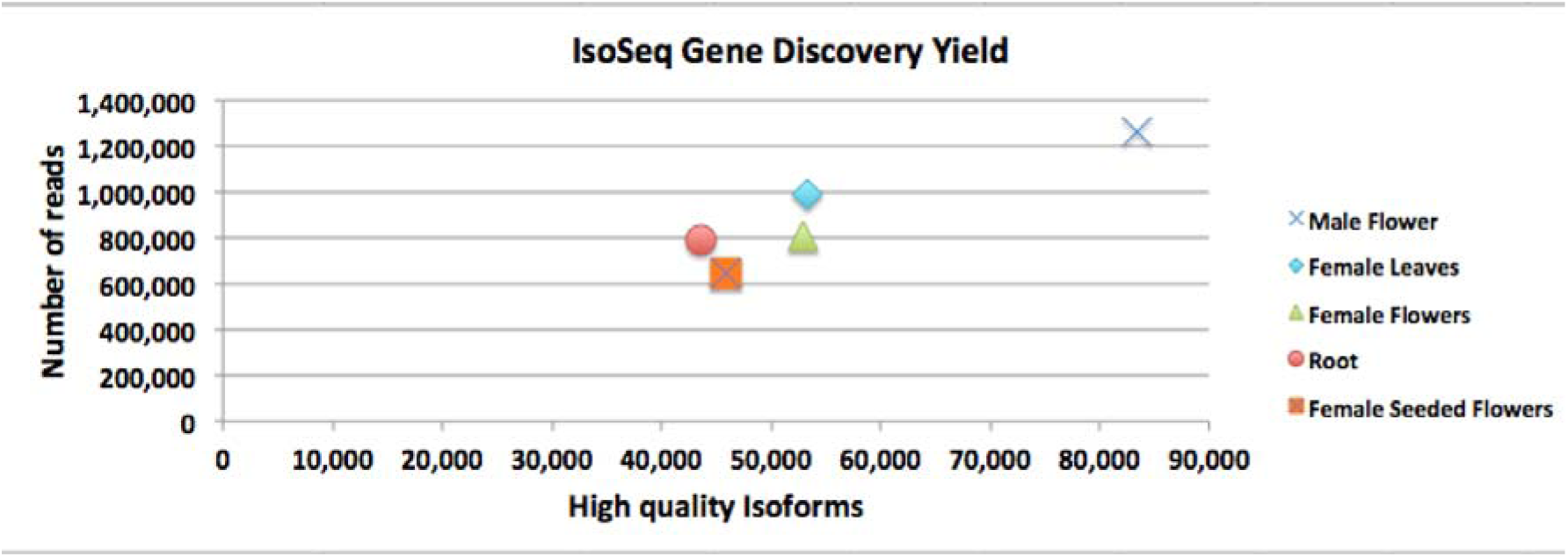
MAKER2 results of Iso-Seq data generated from five tissue types. Isoform discovery is read number-dependent suggesting isoform discovery is not saturated.

**Supplemental Figure S2.**
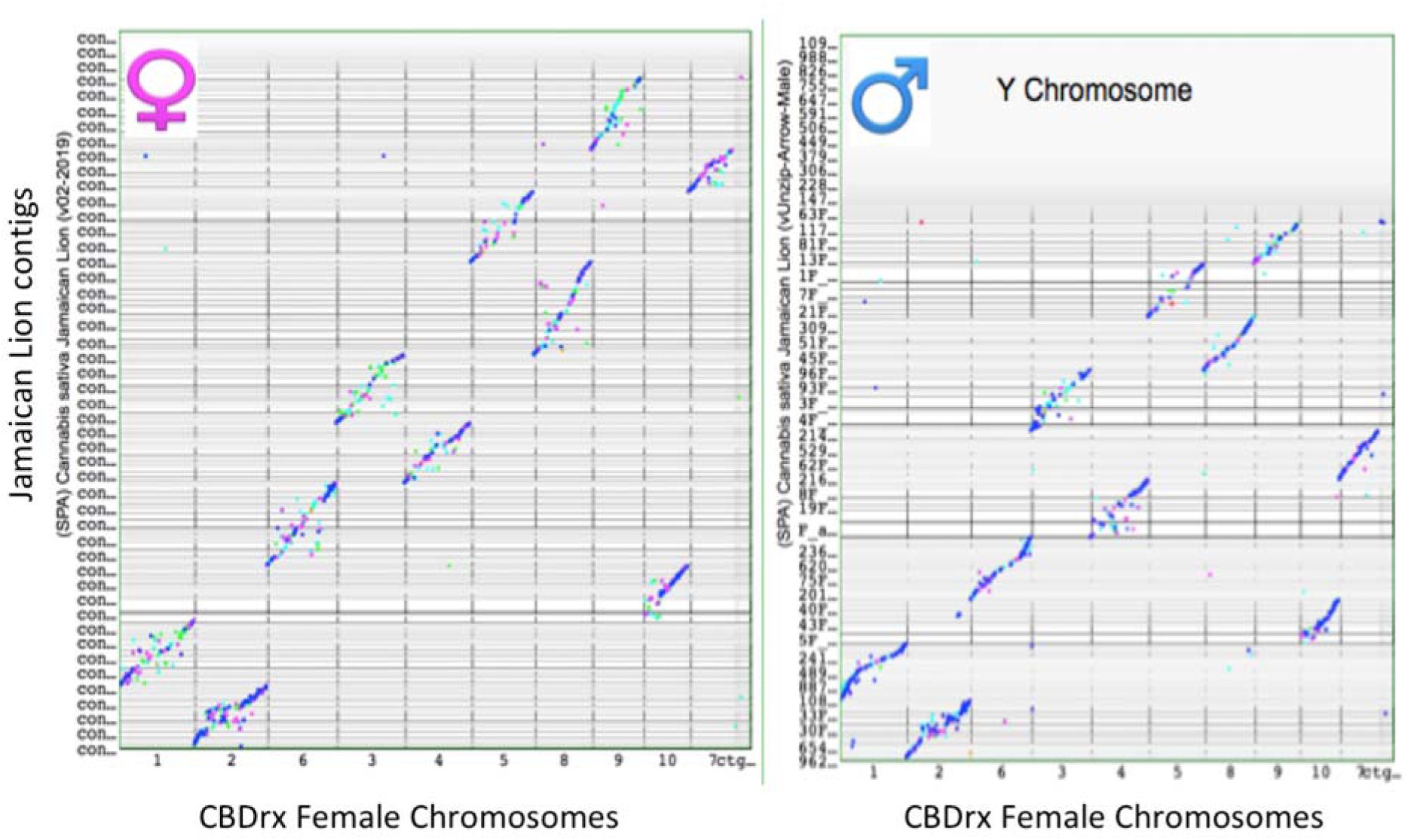
Jamaican Lion parental alignment (female on left, male on right) to CBDrx (Grassa et al. 2018) using SynMap2(Haug-Baltzell et al. 2017). SynMap2 colors dots based on their Ka/Ks values. Ratios of Synonymous (Ks) to Nonsynonmous (Ka) variation can highlight protein coding sequences that have undergone purifying selection. Notice the third largest collection of contigs on Jamaican Lion contigs is chromosome 6 on CBDrx for both Male and Female assemblies. Given the close sizes of cannabis chromosomes and the frequent structural variation in the genome, it is not surprising to see nomenclature discrepancies.

**Supplemental Figure S3:**
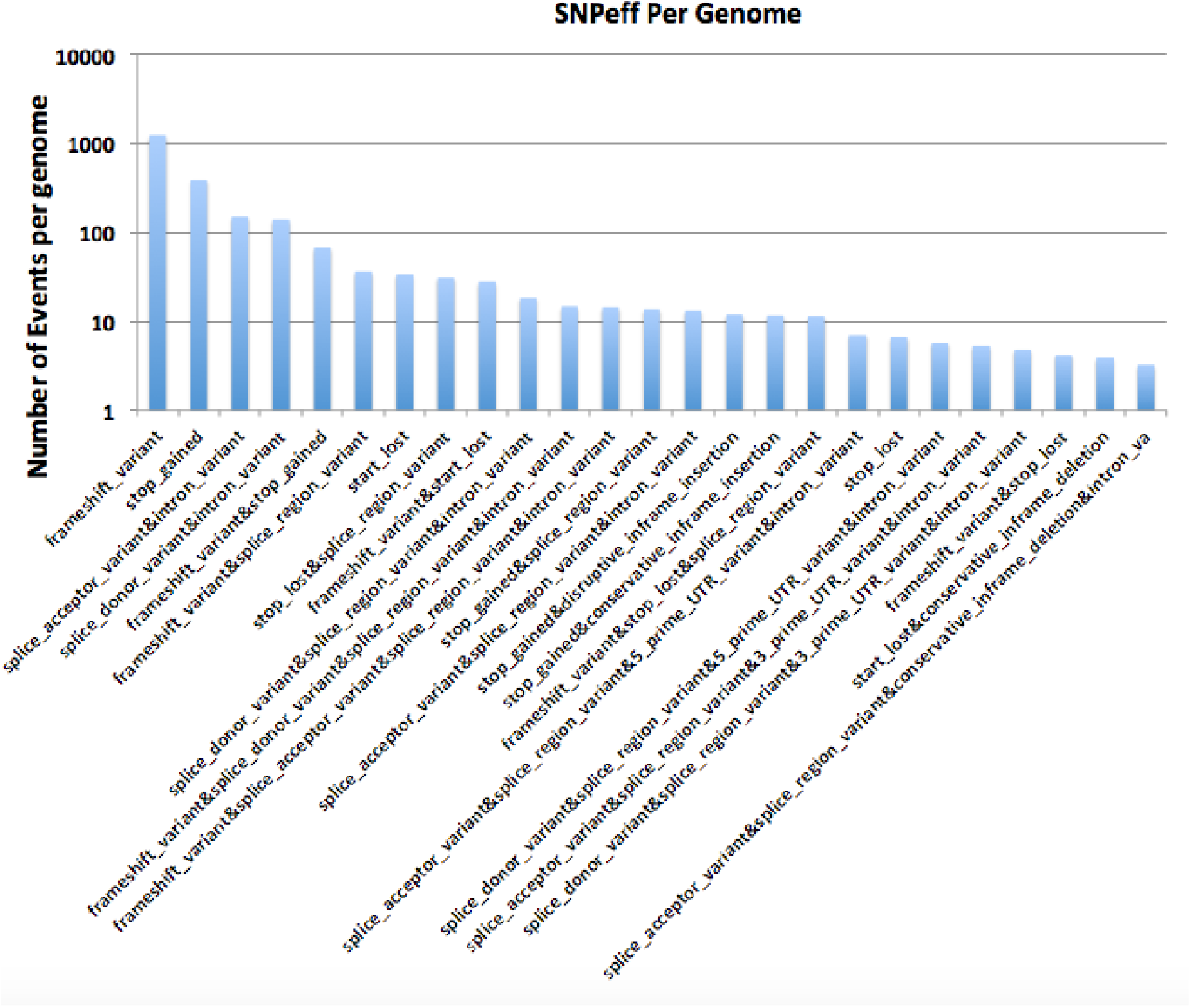
SNPeff calls across the 40 genomes. On average there were over 2,000 high-impact variants per genome (91,144 high impact variants divided by 40 genomes).

**Supplemental Figure S4:**
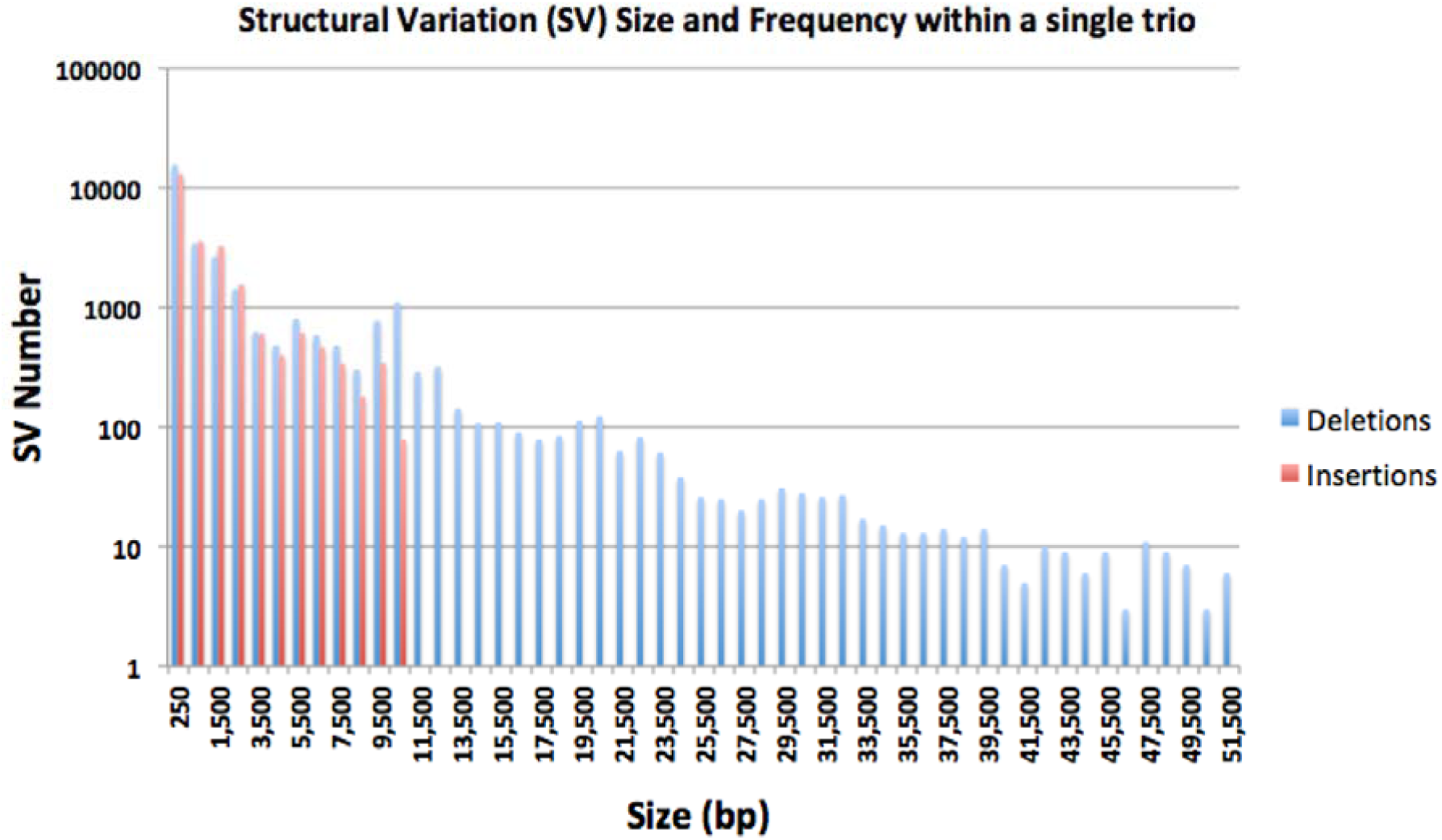
Structural variation detection with PBSV in the Jamaican Lion Trio. Comparison of three genomes (Jamaican Lion mother, Jamaican Lion father, and JL5) results in thousands (y-axis) of structural variations with varying sizes (x-axis). Insertion detection was limited at 10kb detection.

**Supplemental Figure S5.**
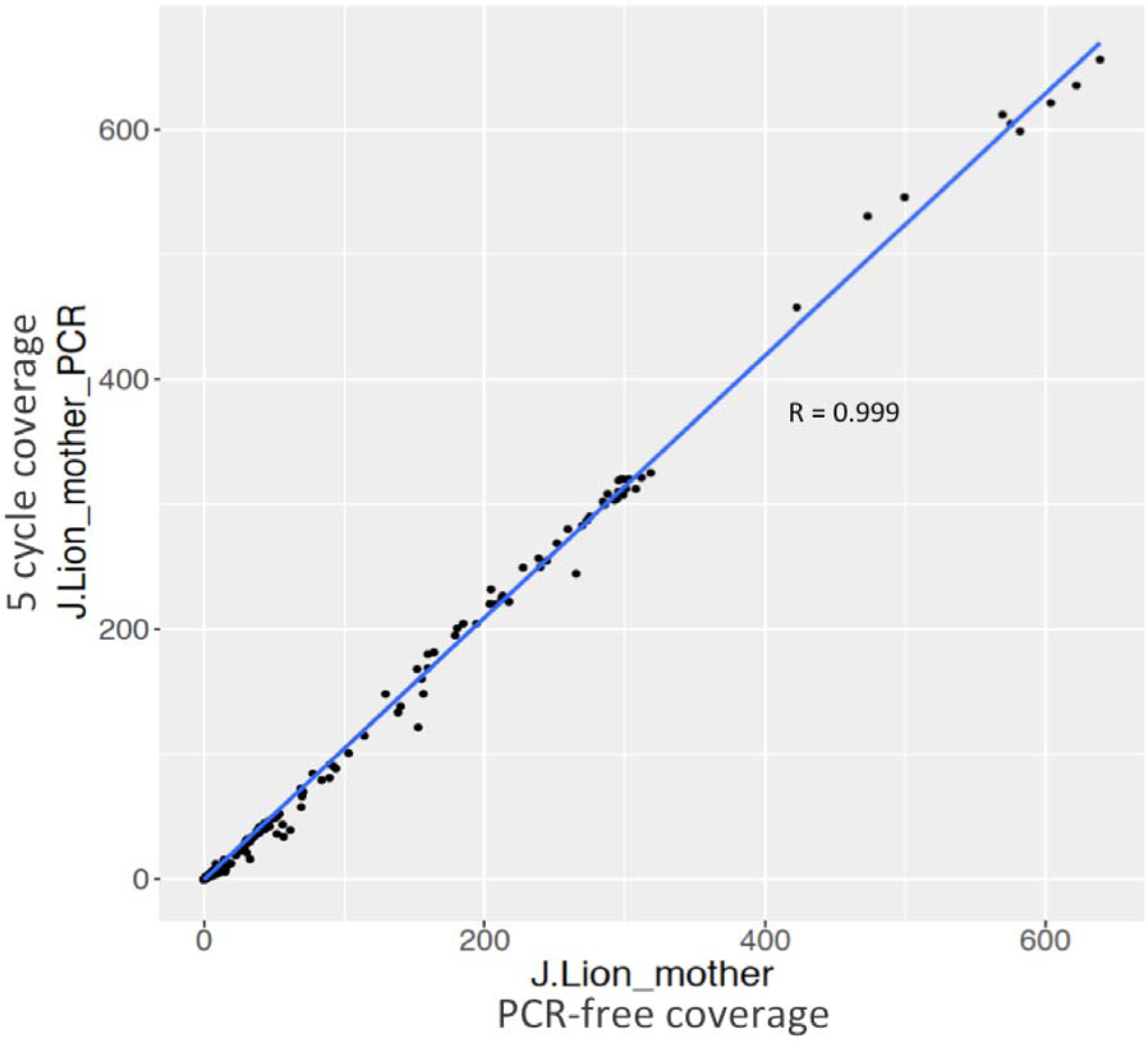
Normalized coverage comparison of 27,644 genes in PCR and PCR-free Jamaican Lion mother libraries including mitochondrial and chloroplast genomes.

**Supplemental Figure S6.**
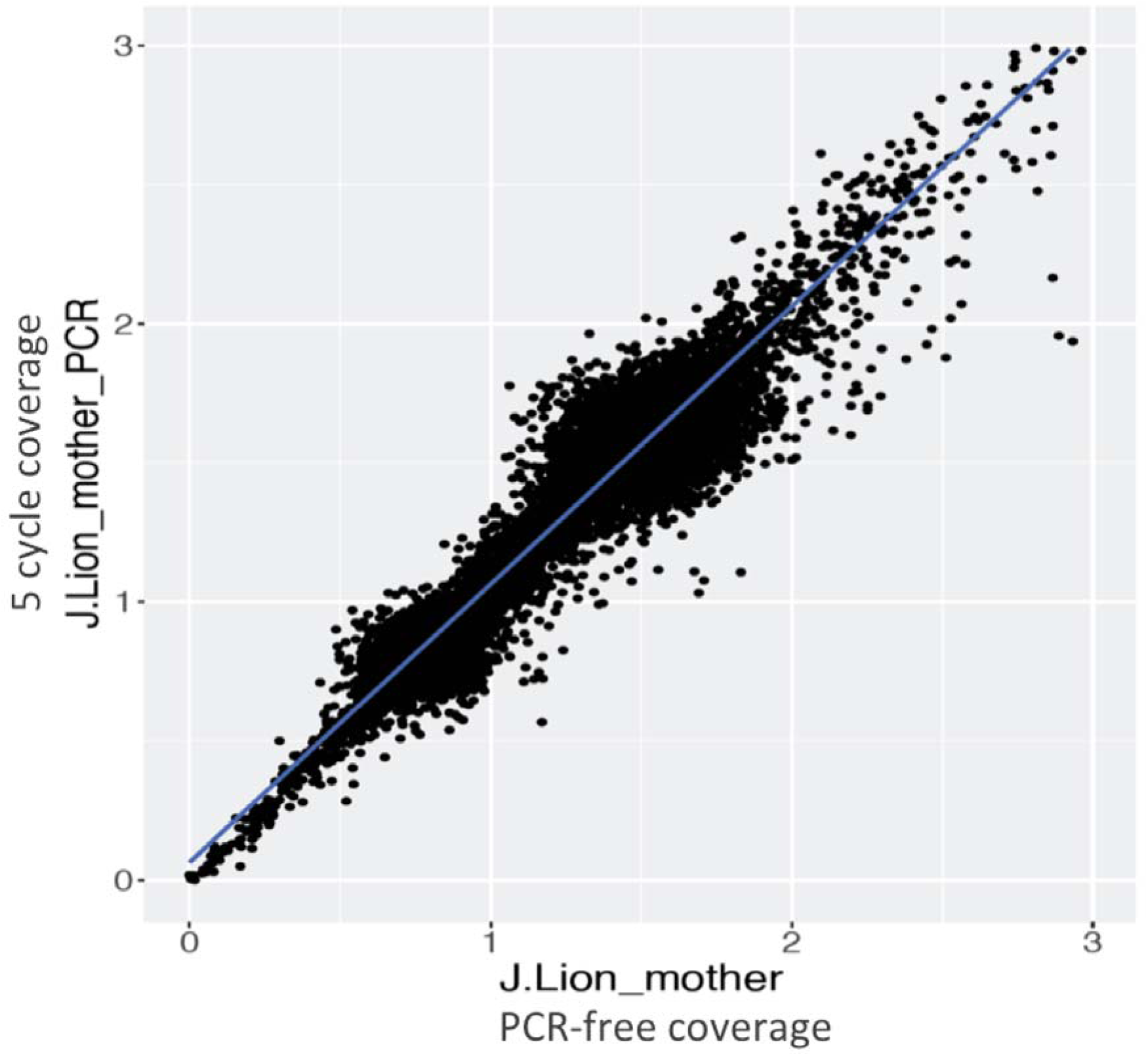
Normalized autosomal gene coverage comparison (without organelle genomes) between PCR-free Illumina sequencing of the maternal Jamaican Lion DNA and 5 cycles of PCR on the same library. Coverage correlation for all genes is very high (r= 0.999).

**Supplemental Figure S7.**
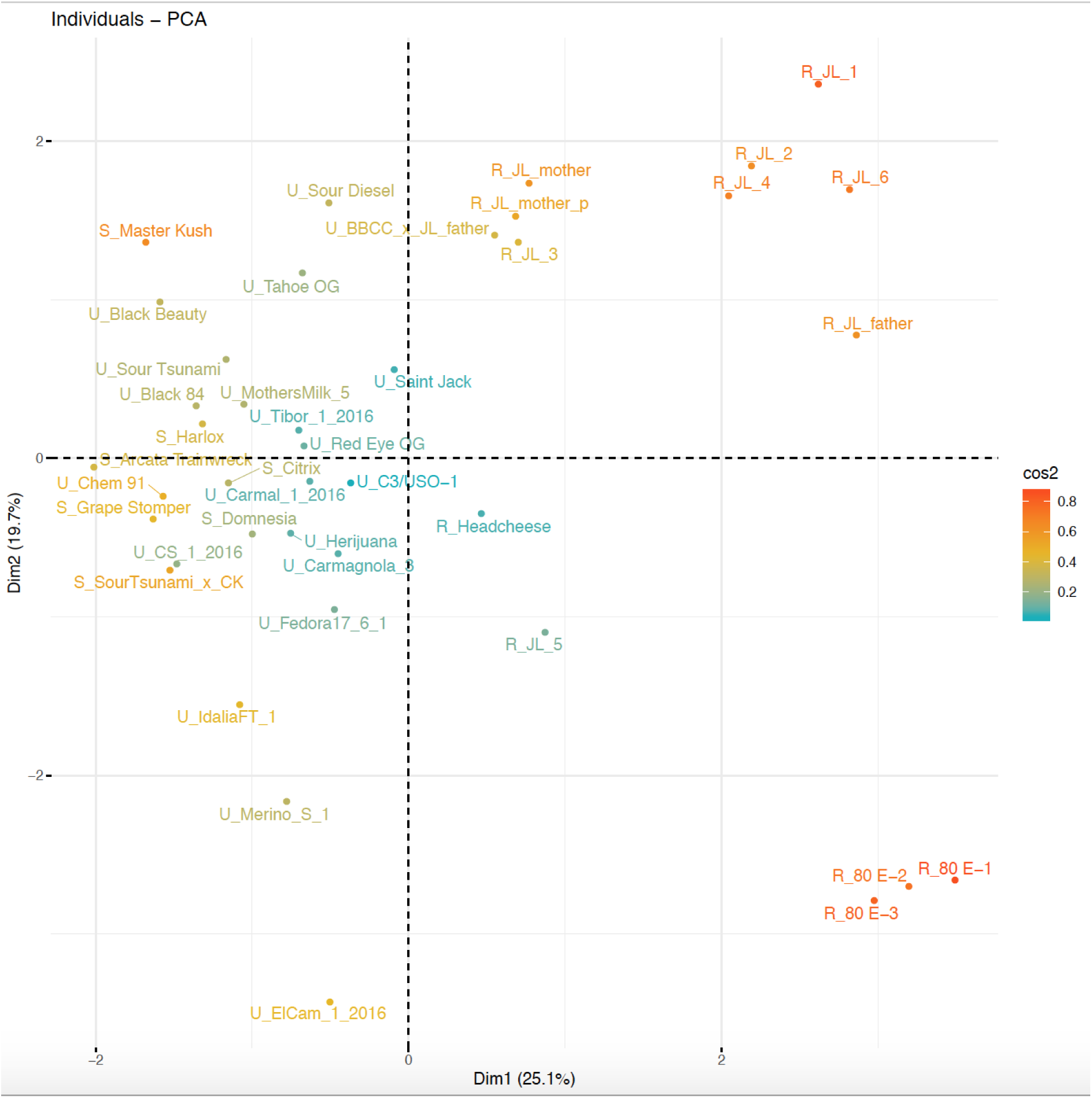
A) Principle component analysis (PCA) of the genomic copy number variation of 23 TLPs, 24 MLOs, and 35 chitinase genes, and pathogen-response genes on the CBCA cluster across 40 genomes. Powdery mildew (PM)- resistant varietals cluster together (bottom right, with prefix “R”). PM-susceptible varietals cluster together (top, with prefix “S”), with unknown PM-resistance are labeled with prefix “U”. Note the paternal Jamaican Lion appears more resistant than the maternal genome. Samples are color-coded based on their cos2 score in the PCA analysis.

**Supplemental Figure S8.**
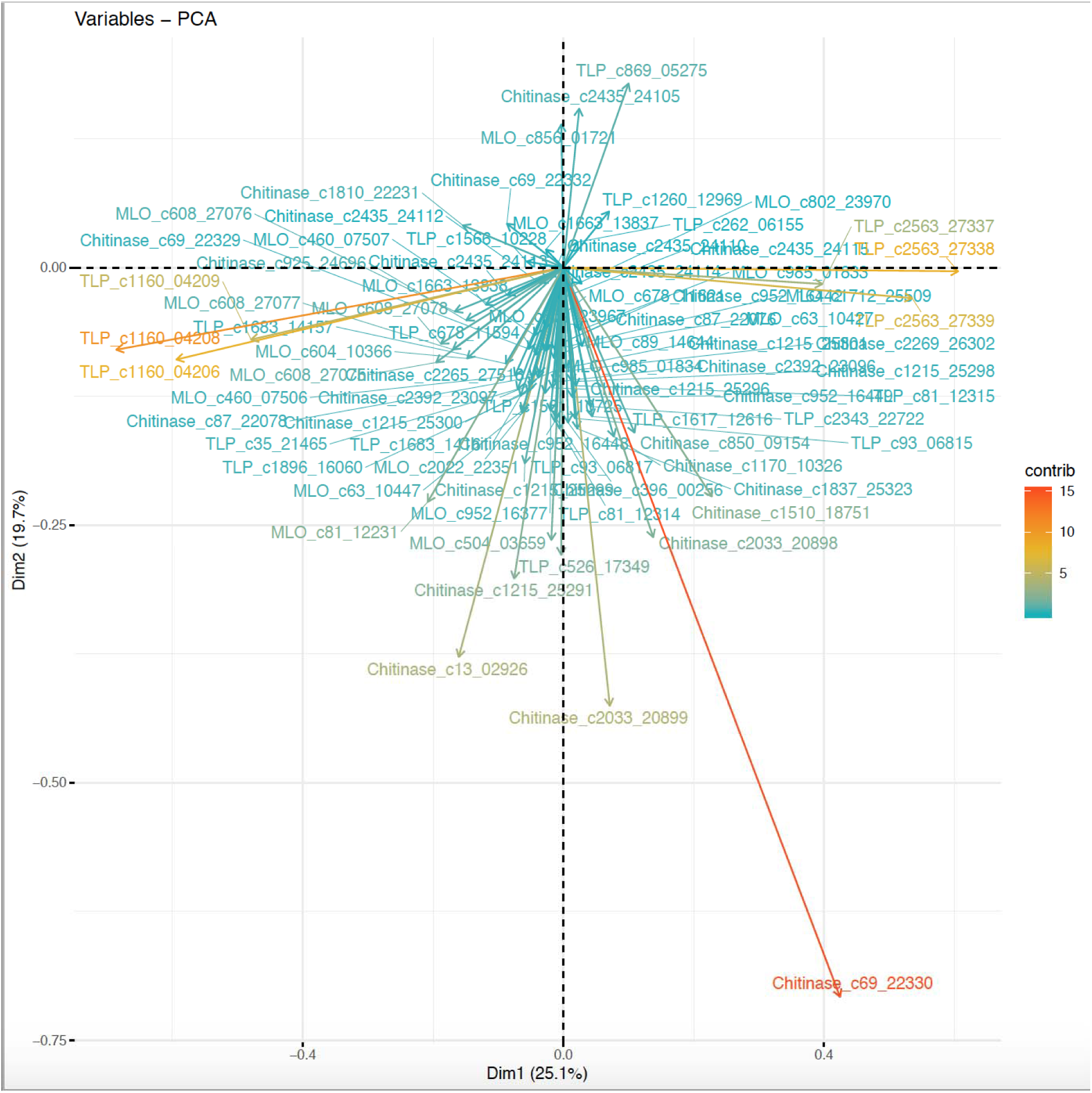
Components in the genomic copy number variation PCA analysis that provide the most segregating contributions are labeled in red and yellow (chitinase c69_22330, TLP_c1160, TLP c2563).

**Supplemental Figure S9.**
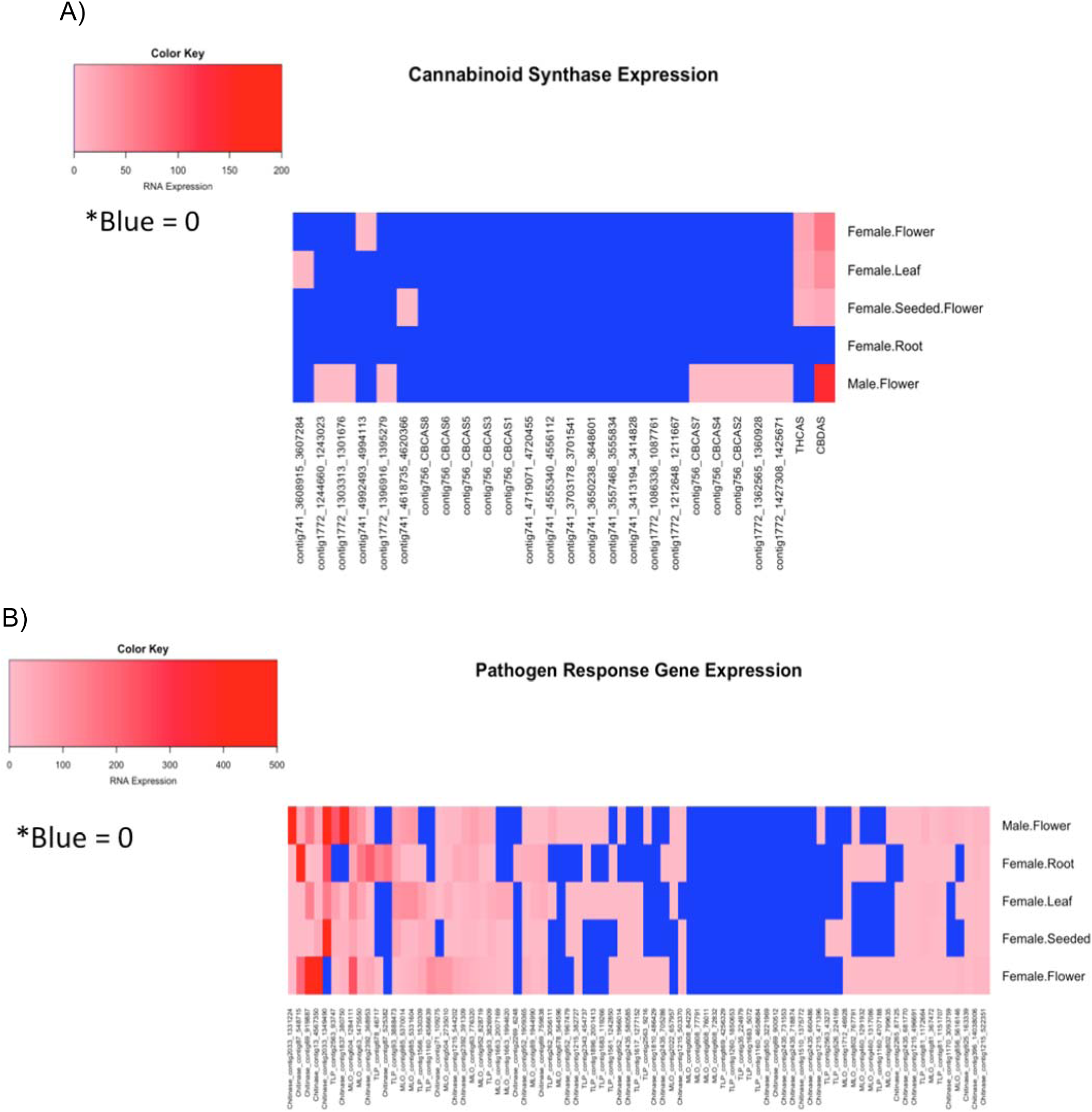
(A)Iso-Seq expression of 26 cannabinoid synthase genes that exist on contig741, contig1772 and contig756. Blue is zero expression, pink to red are increasing transcript levels. (B) Iso-Seq expression levels of 82 pathogen response genes. Zero expression detected is shown in blue. Single transcripts detected are shown in pink and the highest expression levels (1000s) are shown in red. Many of the pathogen response genes with significant segregating power in the PCA analysis are also the most heavily expressed genes (chitinase_c2033, chitinase_c13, chitinase_c69, chitinase_c87, and TLP_2563 or ‘*Cs*TLP1’)

**Supplemental Figure S10:**
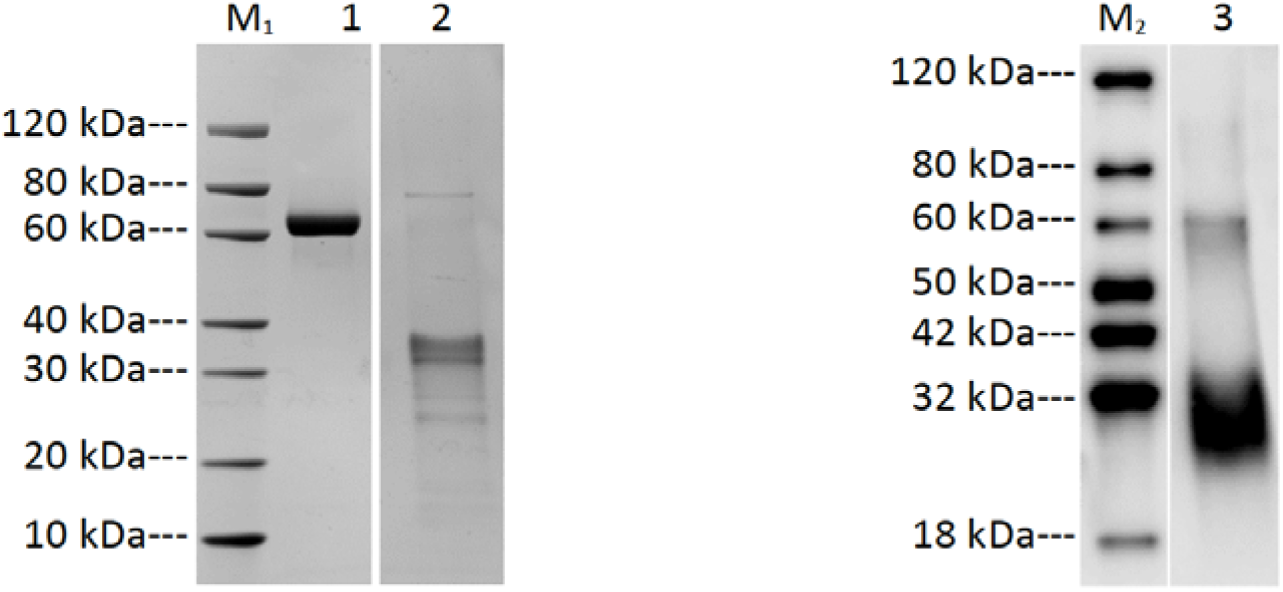
A) SDS-PAGE and B) Western blot analysis of *Cs*TLP1 expression in *E. coli*. Lane M_1_: Protein marker (GenScript, Cat. No. M00516); Lane 1: Bovine serum albumin (2.00 μg); Lane 2: CsTLP1_2563 (reducing condition, 2.00 μg); Lane M_2_: Protein marker (GenScript, Cat. No. M00521); Lane 3: CsTLP1_2563 (reducing condition); primary antibody: mouse-anti-His mAb (GenScript, Cat. No. A00186).

**Supplemental Figure S11.**
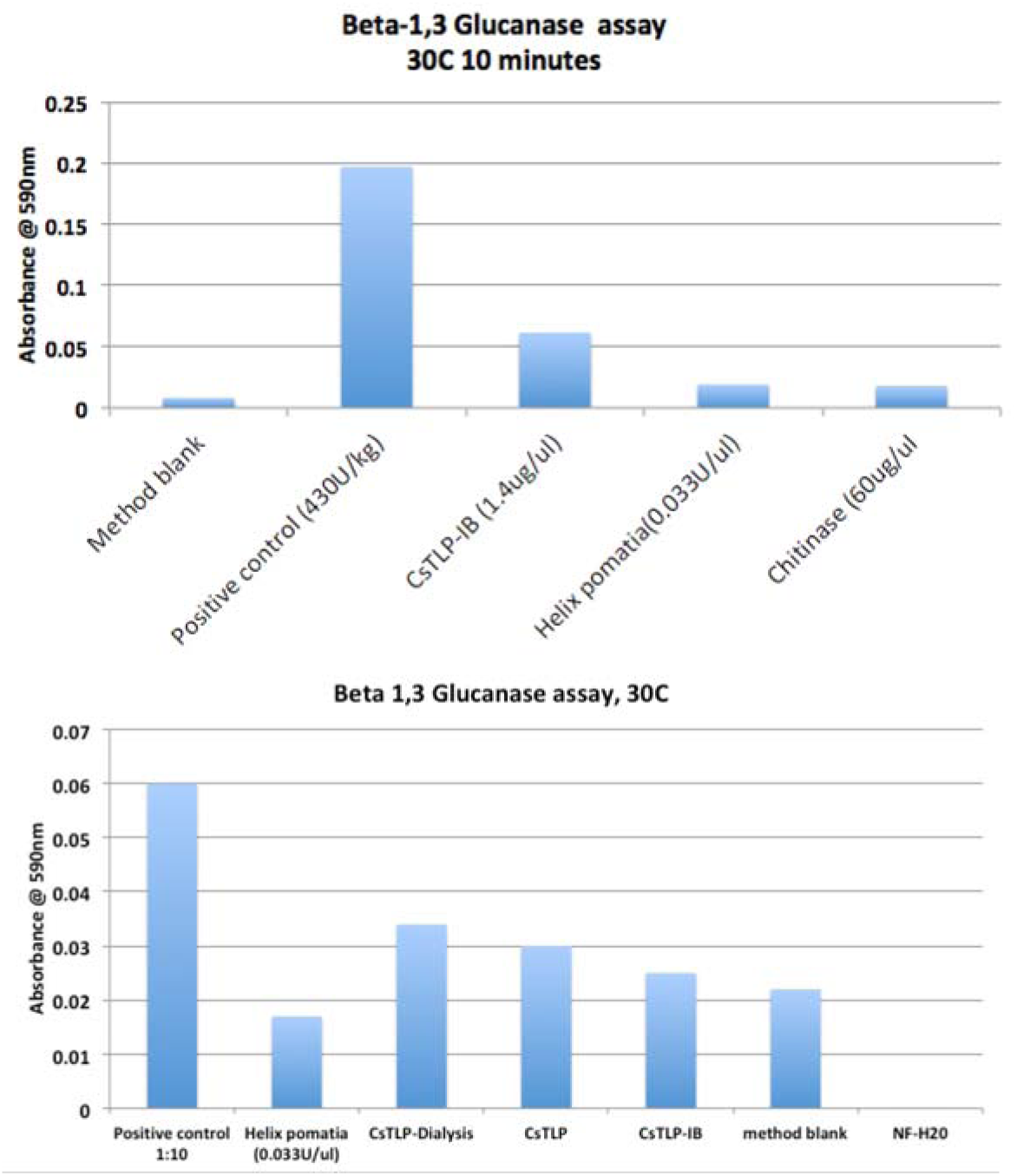
β-glucanase assay (Megazyme) demonstrates activity of the expressed *Cs*TLP1 protein is lower from the inclusion bodies (*Cs*TLP1-IB) fraction than from the cell lysate. Method blanks represent the enzyme activity observed in a few seconds before the precipitation reagent is added. Dialysis of the protein increases *Cs*TLP1 activity implying high-salt buffers may inhibit enzyme activity. Activity is compared to a *Helix pomatia* β-glucanase (Sigma) and a *Trichoderma viride* chitinase (Sigma). The positive control was provided in concentrated form by Megazyme. *H. pomatia* β-glucanase was isolated from a snail stomach with optimal activity at 50 to 55 °C. CsTLP1 has lower activity at 30 °C. Method blank contained the substrate and positive control enzyme but was precipitated with no incubation time.

**Supplemental Figure S12.**
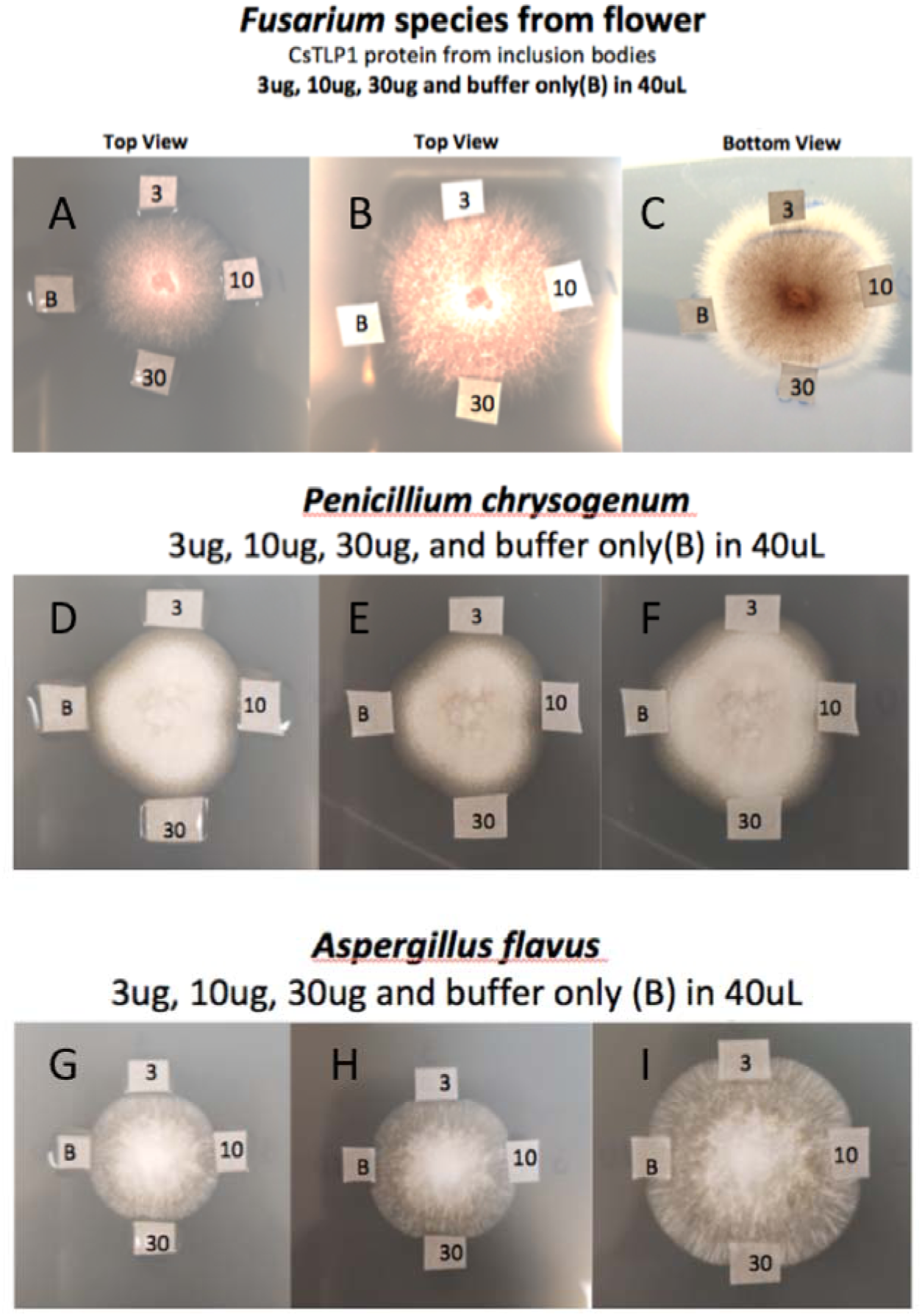
Fungicidal assay of *E. coli* expressing *Cs*TLP1 as described by Misra et al.(Misra et al. 2016) A, B, C) Cannabis derived *Fusarium oxysporum* (qPCR confirmed) grown on potato dextrose agar and treated with varying concentrations of TLP. D, E, F) *Penicillium chrysogenum* (ATCC #18476) and G, H, I) *Aspergillus flavus* (ATCC #16870) and were grown for 48 hours on potato dextrose agar to create a seed colony for exposure to peripheral Whatman paper soaked in thaumatin-like proteins (TLPs) (top: 3 μg in 40 μl^-1^ of *Cs*TLP1; right: 10 μg in 40 μl^-1^ of *Cs*TLP1; bottom: 30 μg in 40 μl^-1^ *Cs*TLP1; left: vehicle control in buffer). *Cs*TLP protein was applied to sterile Whatman paper and placed near organisms growing on potato dextrose agar. Results are shown after approximately 24 h (left), 36h (middle) and 48h (Right). Note, imaging times varied between organisms as each organism grows at a different rate. Limited fungicidal activity is observed with non-dialyzed *Cs*TLP protein. *Cs*TLP1 underwent dialysis to reduce 500mM NaCl concentration for further experiments.

**Supplemental Figure S13.**
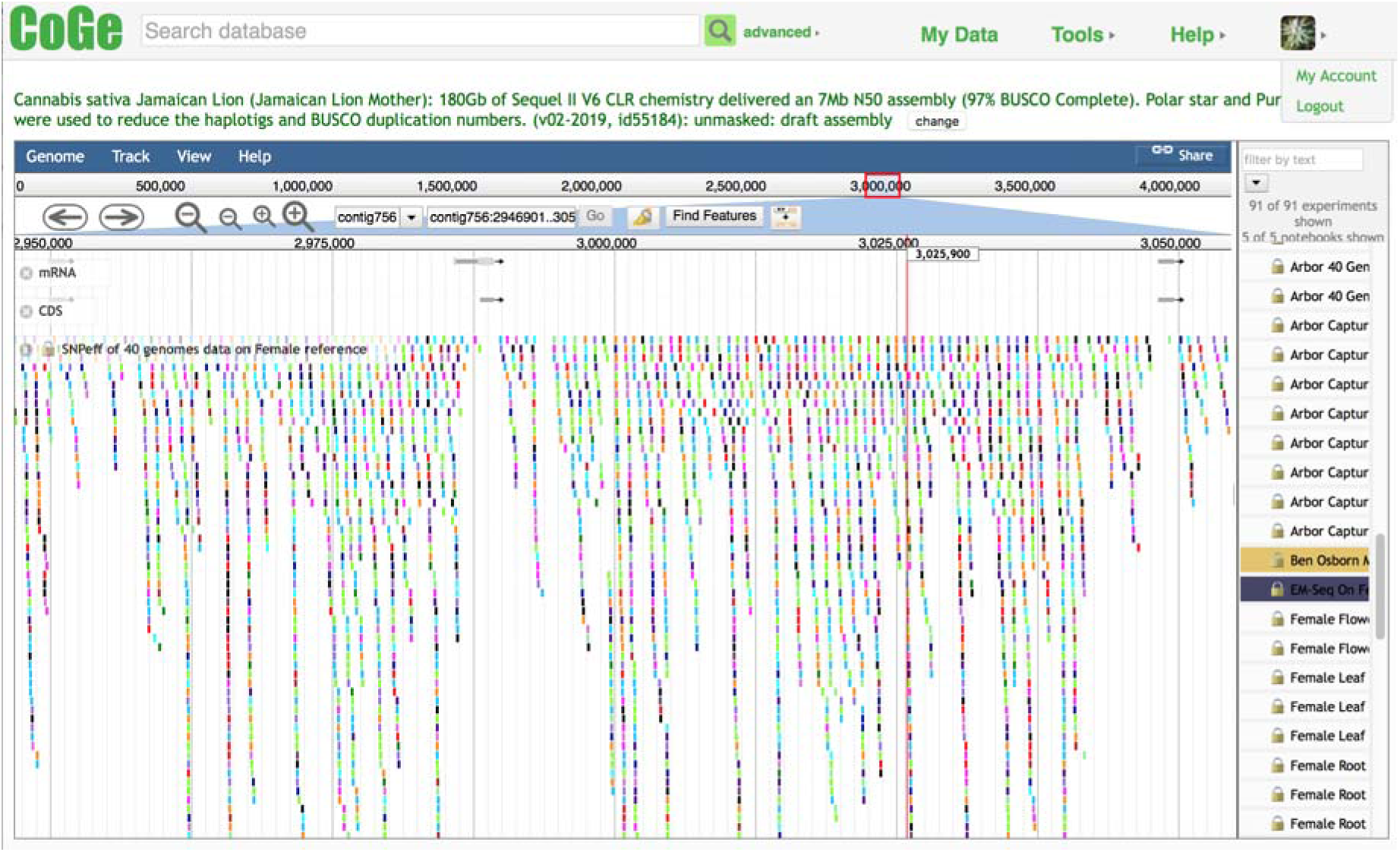
CoGe Genome Browser of 3 of the 8 of the directionally orientated CBCAS genes. These genes are all separated by Copia/LTR sequences. Lower variation in the coding regions can be seen. These references are all available in CoGe at https://genomevo1ution.org/coge/Genomelnfo.pl?gid=55184

**Supplemental Figure S14:**
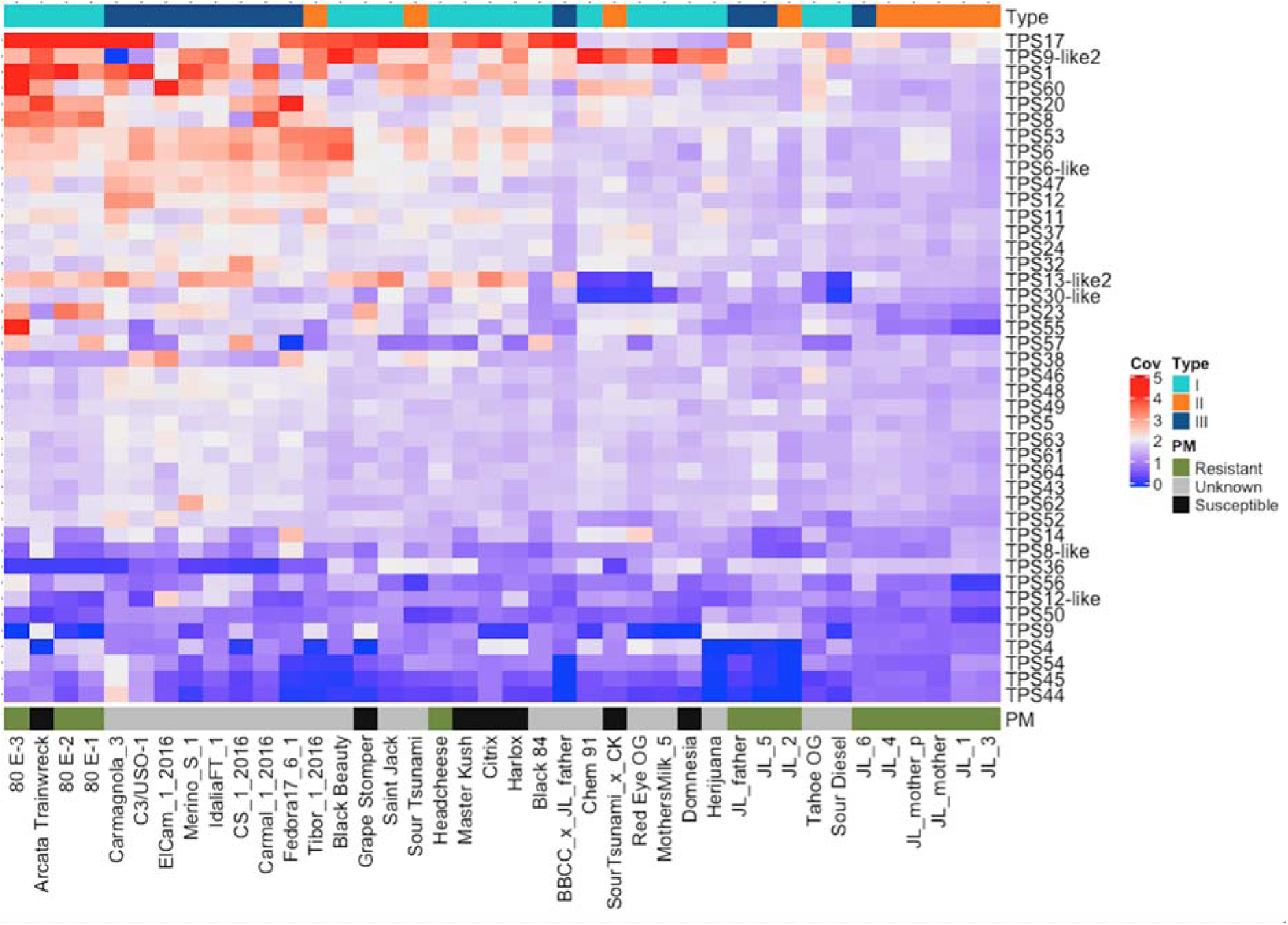
A) A copy number variation (CNV) heatmap of 42 confirmed terpene synthase (TPS) genes. Red is increased copy number and blue represents deleted *Cs*TPS genes.

**Supplemental Figure S15.**
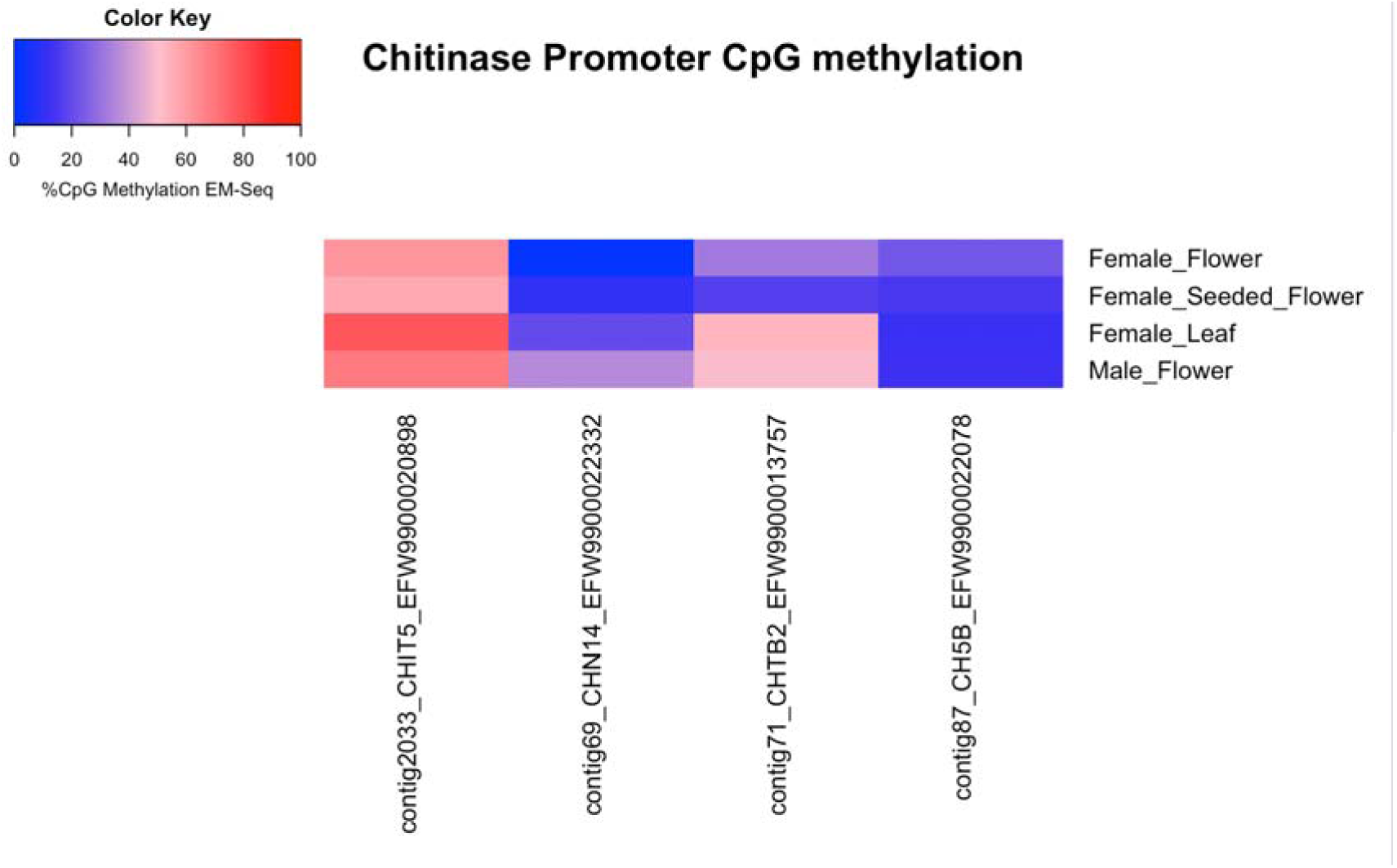
EM-Seq evaluation of CpG methylation 2,000 bases upstream of the Transcriptional Start Sites (TSS) for 4 different sample types (y-axis) of the most differentially expressed genes in the genome. Some of the most differentially expressed genes are 4 chitinases (x-axis).The TSSs that are CpG hypomethylated are shown in blue and the TSSs that are CpG hypermethylated are shown in red. TSSs that are CpG hypomethylated are suggested to be actively transcribed.

**Supplemental Figure S16.**
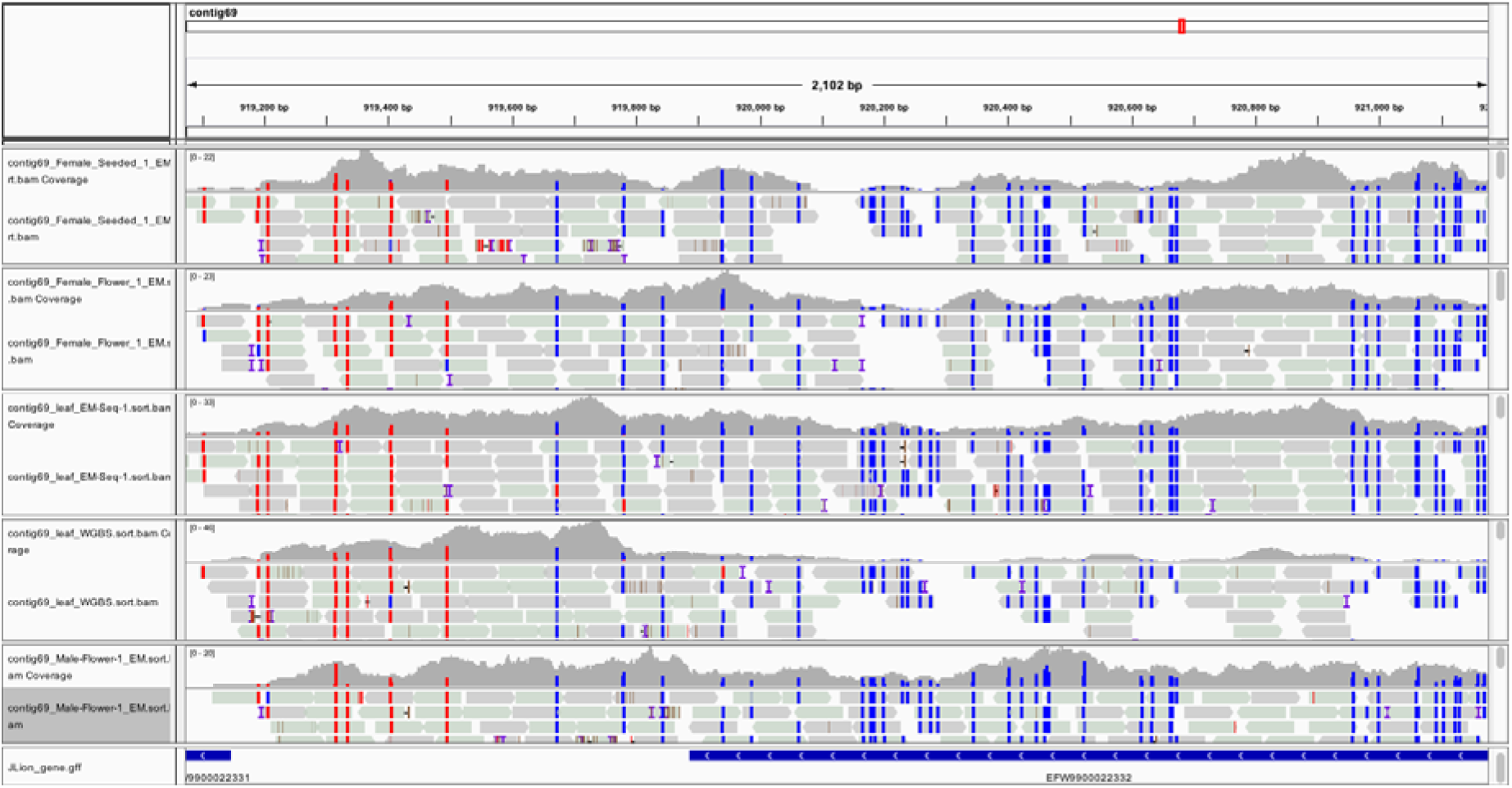
CpG methylation of the CHN14 chitinase gene (contig69) displayed across 4 tissues and one control library. Unmethylated CpGs are shown as blue and methylated bases are shown as red. Four EM-Seq libraries are shown (y-axis). The x-axis is genomic location. Genes with hypomethylated CpGs are suggested to be actively transcribed. IGV_2.4.14 genome browser displays the GFF annotation file as the bottom track.

**Supplemental Figure S17.**
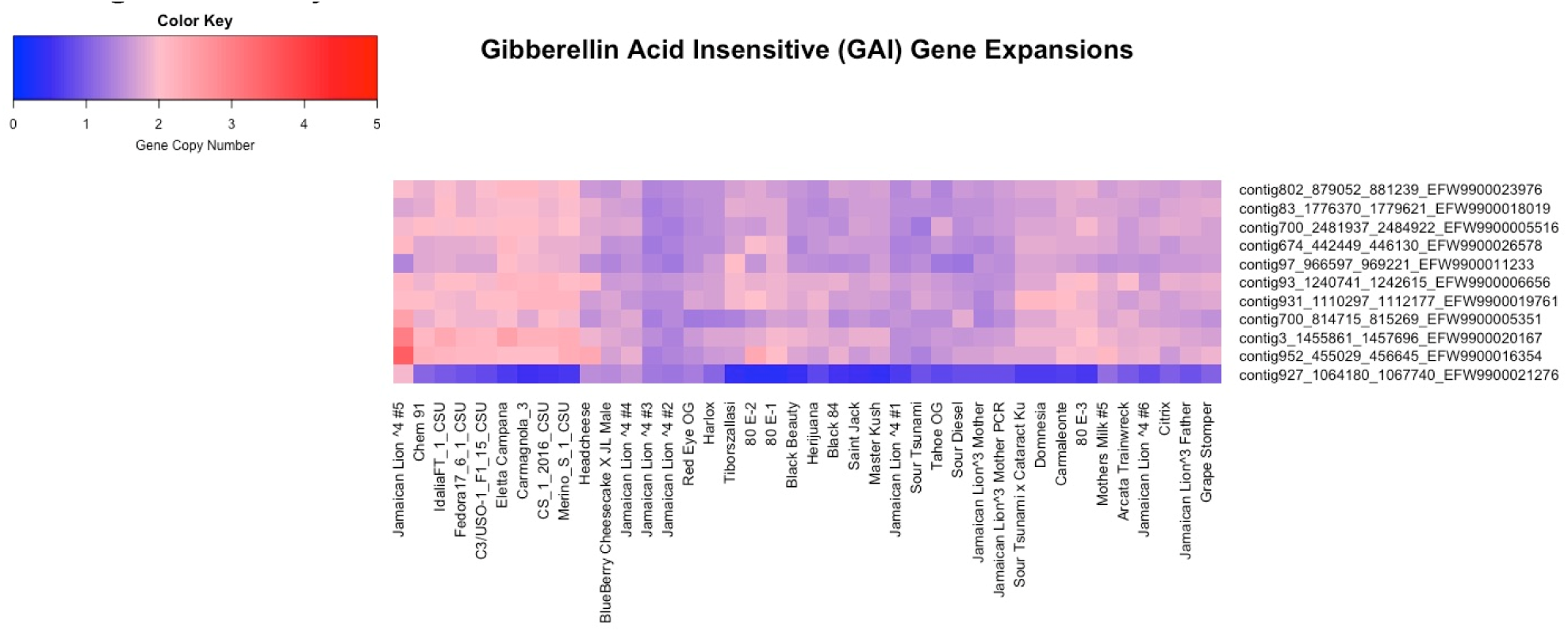
Copy number analysis of 11 Gibberellic Acid lnsenstive (GAI) genes (y-axis) across 40 genomes (x-axis). Two GAI genes are amplified in JL5 (Leftmost and Red) and may be associated with its dwarfism phenotype. Red represents amplified gene copy number and blue signifies deleted genes. Further scrutiny of Gibberellin pathway genetics may lead to a better understanding of cannabis growth and yield.

